# Functional multi-omics reveals genetic and pharmacologic regulation of surface CD38 in multiple myeloma

**DOI:** 10.1101/2021.08.04.455165

**Authors:** Priya Choudhry, Corynn Kasap, Bonell Patiño-Escobar, Olivia Gugliemini, Huimin Geng, Vishesh Sarin, Amrik Kang, Audrey Kishishita, Sham Rampersaud, Letitia Sarah, Yu-Hsiu T. Lin, Neha Paranjape, Poornima Ramkumar, Jonathan C. Patton, Makeba Marcoulis, Donghui Wang, Paul Phojanakong, Veronica Steri, Byron Hann, Benjamin G. Barwick, Martin Kampmann, Arun P. Wiita

## Abstract

CD38 is a surface ectoenzyme expressed at high levels on myeloma plasma cells and is the target for the monoclonal antibodies (mAbs) daratumumab and isatuximab. Pre-treatment CD38 density on tumor cells is an important determinant of mAb efficacy. Several small molecules have been found to increase tumor surface CD38, with the goal of boosting mAb efficacy in a co-treatment strategy. Numerous other CD38-targeting therapeutics are currently in preclinical or clinical development. Here we sought to extend our currently limited insight into CD38 surface expression by using a multi-omics approach. Genome-wide CRISPR-interference screens integrated with patient-centered epigenetic analysis confirmed known regulators of CD38, such as RARA, while revealing XBP1 and SPI1 as other key transcription factors governing surface CD38 levels. CD38 knockdown followed by cell surface proteomics demonstrated no significant remodeling of the myeloma “surfaceome” after genetically-induced loss of this antigen. Integrated transcriptome and surface proteome data confirmed high specificity of all-trans retinoic acid in upregulating CD38 in contrast to broader effects of azacytidine and panobinostat. Finally, unbiased phosphoproteomics identified inhibition of MAP kinase pathway signaling in tumor cells after daratumumab treatment. Our work provides a resource to design strategies to enhance efficacy of CD38-targeting immunotherapies in myeloma.

## Introduction

Harnessing the immune system to treat myeloma has rapidly become the most exciting therapeutic frontier in this disease. The first such immunotherapy agent to achieve United States Food and Drug Administration (FDA) approval was the monoclonal antibody (mAb) daratumumab^1^. Daratumumab targets CD38, a cell surface ectoenzyme highly expressed on myeloma plasma cells. Daratumumab is currently FDA-approved for use as either monotherapy or combination therapy in the relapsed/refractory setting, or front-line therapy in combination with other small molecule agents^1^. A second mAb targeting CD38, isatuximab, was also recently approved for relapsed/refractory myeloma; at least 15 additional CD38-targeting agents are in development^2^. Extensive and encouraging clinical data has already been obtained with daratumumab, though resistance appears to inevitably occur^3,4^. Biologically, this process appears to be quite complex, with determinants of resistance ranging from alteration of surface antigens on tumor cells^4–6^ to dysfunction of the tumor immune microenvironment^7,8^. While it remains unclear whether CD38 downregulation on tumor cells after mAb treatment is a marker of resistance^5,9^, or, instead, successful therapy^10^, compelling preclinical and clinical data suggests that CD38 surface antigen density prior to treatment strongly correlates with mAb efficacy^5,11^.

This latter observation has led to numerous efforts to identify small molecules which can increase tumor surface antigen density of CD38, representing potential co-treatments with CD38-targeting mAbs. The first such example of a CD38-boosting small molecule was all-trans retinoic acid (ATRA)^12^. Subsequent studies identified the pan-histone deacetylase (HDAC) inhibitor panobinostat^13^, the thalidomide analog lenalidomide^14^, the Janus kinase (JAK) inhibitor ruxolitinib^15^, and the DNA methyltransferase inhibitor azacytidine^16^ as agents that could lead to myeloma surface CD38 increase. A clinical trial combining ATRA with daratumumab has led to encouraging outcomes in patients previously refractory to daratumumab^17^.

While these published strategies suggest ways to improve CD38 mAb outcomes, they also leave many questions unanswered. Most notably, we do not yet have a broad global sense of the transcriptional or post-transcriptional networks that most strongly impact CD38 expression. Bi- and tri-specific antibodies^18^ and chimeric antigen receptor (CAR) T cells^19^ targeting CD38 are also in clinical development. As seen for similar modalities against other targets^20^, efficacy of these novel agents, in addition to mAbs, is likely to also be impacted by CD38 antigen density on tumor cells. Furthermore, prior studies showed that CD38 downregulation after daratumumab treatment was accompanied by increases in the complement inhibitors CD55 and CD59 (ref.^5^). Are there other features of myeloma surface remodeling driven by CD38 downregulation? And, for the small molecules noted above, it is unknown how they more generally impact the makeup of the myeloma cell surface proteome beyond CD38. The tumor cell surface not only harbors opportunities for immunotherapeutic targeting but also is the interface for communication between tumor and microenvironment, potentially leading to other alterations in myeloma biology after changes in surface proteomic profile. To address these questions, here we have taken advantage of CRISPR interference-based functional genomic screens, cell surface proteomics, epigenetic analyses, and phosphoproteomics to provide a multi-omic perspective on CD38 regulation and tumor cell consequences of targeting CD38 in myeloma.

## Methods

### CRISPR interference screening and hit validation

Genome-wide CRISPRi screening was performed as described previously^21^. Briefly, RPMI-8226 cells stably expressing dCas9-KRAB were transduced with a genome-wide library comprised 5 sgRNA/protein coding gene. After 14 days cells were stained for surface CD38 and flow-sorted to enrich for populations of cells expressing low or high cell surface levels of CD38. Cell populations were then processed for next-generation sequencing as previously described^22^ and sequenced on a HiSeq-4000 (Illumina). Reads were analyzed by using the MAGeCK pipeline as previously described^23^. Further validation was performed by knockdown with individual sgRNA’s extracted from the genome-wide library with conformation by flow cytometry or Western blotting. Antibody-dependent cytotoxicity assays were performed using NK92-CD16 cells as described previously^16^. Additional details in Supplementary Methods.

### Epigenetic analysis and machine learning for CD38 transcriptional regulation

Publicly available ATAC-seq data from primary myeloma samples (ref.^24^) was analyzed with the Homer tool findPeaks. Motif binding in the identified ATAC peak regions was called with PROMO tool^25^. Newly-diagnosed patient tumor RNA-seq data in the Multiple Myeloma Research Foundation CoMMpass trial (MMRF; research.themmrf.org) was used to correlate expression of predicted transcription factors with *CD38* expression. To build a predictive model for *CD38* expression as a function of transcription factor expression, we developed an XGBoost (Extreme Gradient Boosting) model with randomized search with cross validation to find optimal parameters. 80% of CoMMpass data was used for training and the remainder for model testing. Additional details in Supplementary Methods.

### Cell surface proteomics and phosphoproteomics

Cell surface proteomics was performed using an adapted version of the *N*-glycoprotein Cell Surface Capture^26^ method, as we have described previously^27^. Unbiased phosphoproteomics was performed using immobilized metal affinity (IMAC) chromatography using methods described previously^28^. All samples were analyzed on a Thermo Q-Exactive Plus mass spectrometer with data processing in MaxQuant^29^. Additional details in Supplementary Methods.

## Results

### A CRISPR interference-based screen reveals regulators of CD38 surface expression

We first sought to use an unbiased approach to identify regulators of surface CD38 in myeloma tumor cells. We specifically employed genome-wide screening with CRISPR interference (CRISPRi), which leads to much higher specificity of knockdown than shRNA while avoiding potential toxicity of double-strand breakage with CRISPR deletion^30^. We recently used this approach to characterize regulators of surface B-cell Maturation Antigen (BCMA) in myeloma^21^. Here, we employed an RPMI-8226 cell line with the dCas9-KRAB machinery, required for CRISPRi, as described previously^21^. We confirmed that this RPMI-8226 cell line robustly expressed CD38 (**Supp. Fig. 1A**).

The genome-wide screen was performed as shown in **Fig. 1A**. Briefly, RPMI-8226 cells were transduced with a pooled genome-wide sgRNA library. After 14 days the cells were then stained with fluorescently-labeled anti-CD38 antibody and flow sorted into low- and high-CD38 populations. Frequencies of cells expressing each sgRNA was quantified using next generation sequencing. As an important positive control, increasing confidence in the screen results, we first noted that knockdown of *CD38* itself strongly decreased surface CD38 expression (**Fig. 1B**). On the other hand, several dozen genes, when repressed, did indeed lead to increased surface CD38 (right side of volcano plot in **Fig. 1B; Supp. Table 1**). As another positive control, one of these top hits included *RARA*, whose degradation is catalyzed by ATRA treatment to drive CD38 increase^12^.

**Figure 1.**
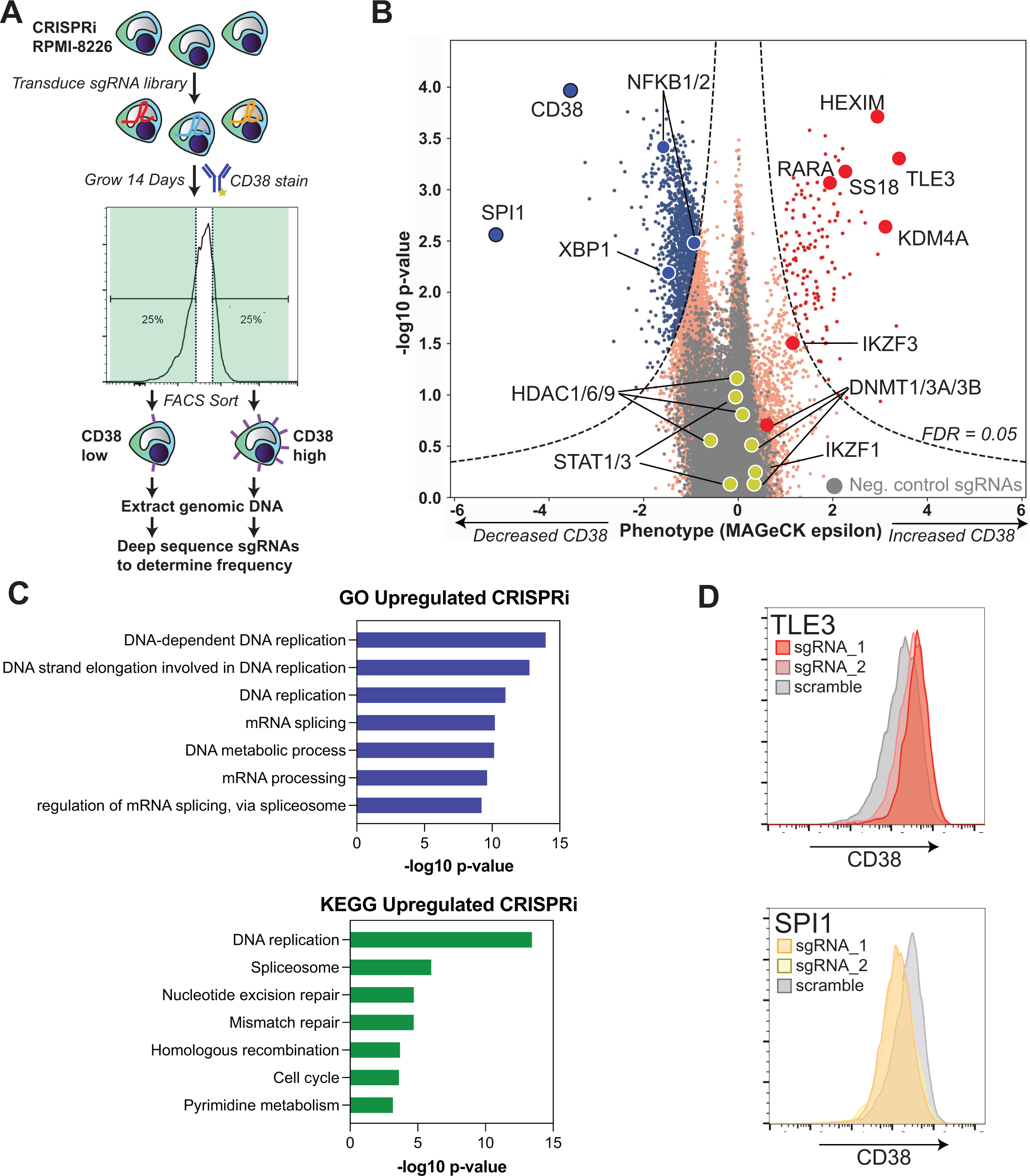
CRISPR interference (CRISPRi) screening reveals genetic determinants of surface CD38 regulation. **A.** Schematic of CRISPRi screen design. **B.** Results of CRISPRi screen demonstrating genes that, when knocked down, regulate surface CD38 in RPMI-8226 cells. X-axis indicates phenotype (epsilon) from MAGeCK^55^ statistical analysis. Dashed line indicates cutoff for significant change at False Discovery Rate (FDR) < 0.05. Genes of interest are specifically labeled. 4,000 negative control non-targeting sgRNAs are in grey. **C.** Gene Ontology (GO) Biological Process and Kyoto Encyclopedia of Genes and Genomes (KEGG) analysis of all genes that when knocked down lead to significant CD38 upregulation. **D.** Follow-up flow cytometry validation of CRISPRi screen hits using two individual sgRNAs per gene demonstrates *TLE3* knockdown drives increased CD38, while *SPI1* knockdown leads to CD38 decrease.

To find pathways that may be useful for pharmacologic targeting, we first applied Gene Ontology (GO) and Kyoto Encyclopedia of Genes and Genomes (KEGG) analysis to the list of genes that, when inhibited, significantly increased CD38 (**Fig. 1C**). We were intrigued to find that many of the strongest effects appeared to be driven by transcriptional or other epigenetic factors. These specifically included pathways such as “DNA replication”, “mRNA processing”, “DNA-templated transcription”, and “spliceosome”.

We considered whether any hits associated with these pathways may be “druggable”, with the goal of expanding our repertoire of small molecules that enhance surface CD38 in myeloma. *SS18*, a component of the BAF (BRG1/BRM associated factor) chromatin remodeling complex, scored highly as a hit. However, treatment with the proposed BAF inhibitor caffeic acid phenol ester (CAPE)^31^ did not lead to consistent increases in surface CD38 (**Supp. Fig. 1B**). Similarly, the lysine demethylase *KDM4A* was a prominent hit, but treatment with the inhibitory metabolite (*R*)-2-hydroxyglutarate^32^ also had no effect (**Supp. Fig. 1B**).

The strongest hits for genes whose knockdown increased surface CD38 were two transcription factors, *HEXIM1* and *TLE3*. Validation studies using individual sgRNA knockdown confirmed increased surface CD38 (**Fig. 1D**, **Fig. 2A** and **Supp. Fig. 2A**), as well as functional impact in NK cell antibody-dependent cellular cytotoxicity (ADCC) assays with daratumumab (**Fig. 2B**). However, these proteins are known to be widespread negative regulators of transcription^33,34^, suggesting little scope for specific therapeutic targeting at the *CD38* locus.

**Figure 2.**
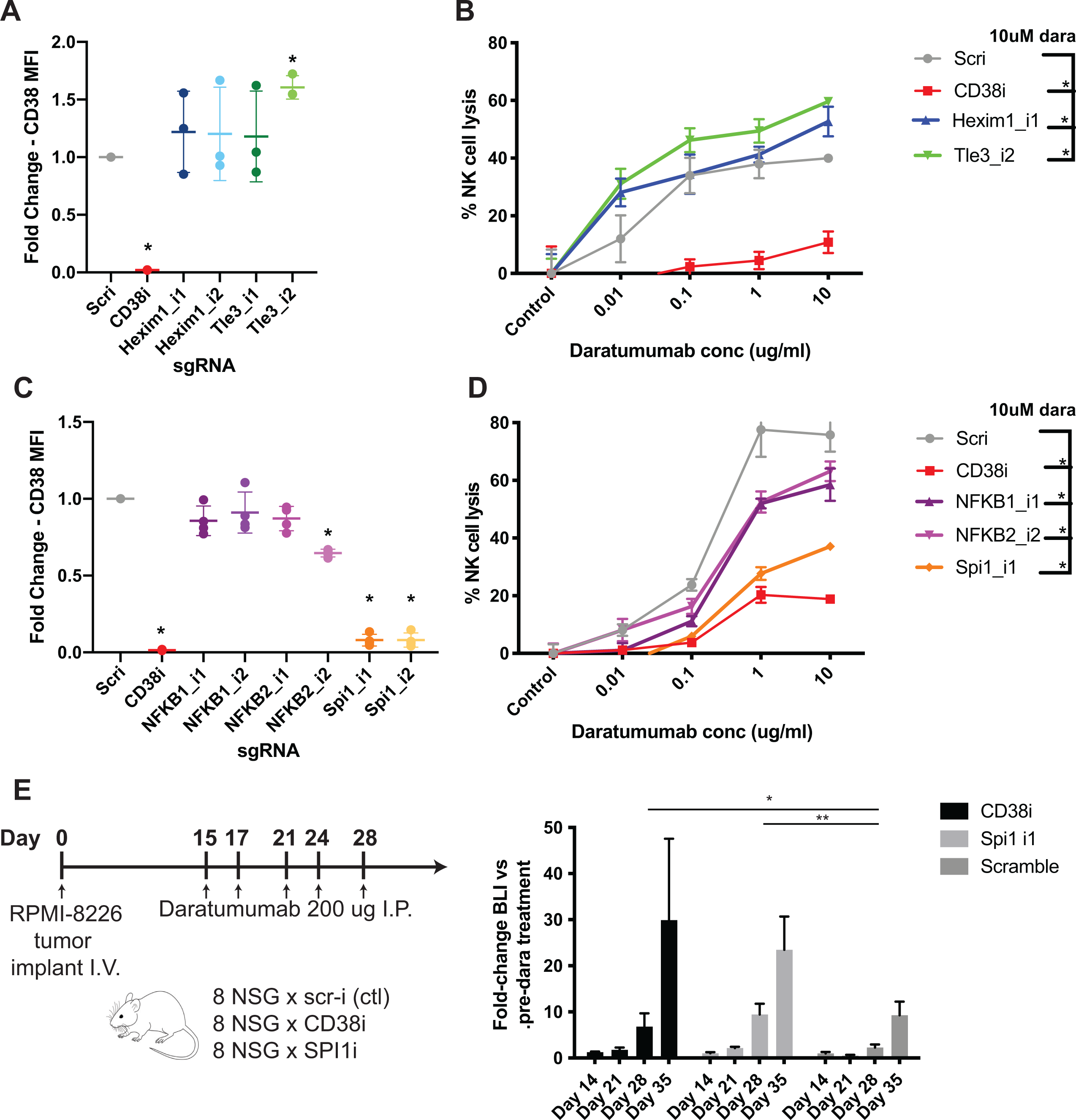
Validation of CRISPRi screen hits as functionally impacting daratumumab efficacy. **A.** Knockdown of HEXIM1 and TLE3 with two independent sgRNA’s/gene (AMO1 myeloma cells, *n* = 3) followed by flow cytometry shows significant surface CD38 increase with TLE3_i2 sgRNA and trend toward increased CD38 with HEXIM1_i1 sgRNA. Scri = non-targeting control sgRNA. **B.** Results from ADCC assays with AMO1 cells stably expressing the noted sgRNA’s and incubated with the indicated concentration of daratumumab or isotype control antibody (1:20 myeloma:NK ratio, 20 hours, *n* = 2). The percent lysis by ADCC was calculated using the following formula : % Lysis = (signal in presence of daratumumab – signal in presence of IgG1 control antibody) x100 / signal in presence of IgG1 control antibody. At 10 μM daratumumab, both HEXIM1 and TLE3 knockdown led to significant increase in ADCC. **C.** Similar to A., sgRNA knockdown of *NFKB1*, *NFKB2*, and *SPI1* with fold-change in CD38 by flow cytometry (RPMI-8226 cells, *n* = 3). **D.** Similar to B., knockdown with the most effective sgRNA for each gene show significant decreases in NK-cell ADCC at 10 μM daratumumab in the RMPI-8266 cells (*n* = 3). **E.** *In vivo* validation of *SPI1* knockdown driving daratumumab resistance. NOD *scid* gamma mice were I.V. implanted with CRISPRi RPMI-8226 cells stably expressing both luciferase and noted sgRNA, then treated with 200 μg daratumumab on the noted schedule. Bioluminescence imaging measurement of tumor burden demonstrates significantly increased fold-change in tumor burden (normalized to pre-daratumumab intensity) with either CD38 or SPI1 knockdown compared to scramble sgRNA. For **A**-**E**, \**p* < 0.05, \*\**p* < 0.01 by two-tailed *t*-test.

We were surprised that other targets proposed to increase CD38 expression after pharmacologic inhibition, such as HDACs^3^, or catalyzed degradation, such as IKZF1/3 (ref.^25^), did not appear as prominent hits (**Fig. 1B**). However, this result may reflect a limitation of functional genomic screens. A pharmacologic agent may inhibit multiple members of a protein class to drive a phenotype, whereas, with single gene knockdown, functional redundancy may prevent this phenotype from appearing^35^ (i.e., multiple HDACs may need to ablated at once, or both IKZF1 and IKZF3 simultaneously, to drive increased CD38). We speculate this is the case with DNA methyltransferases (DNMTs). We previously showed that treatment with the DNMT inhibitor azacytidine, which promotes degradation of all cellular DNMTs^36^, could robustly increase surface CD38 (ref.^16^). Here, however, we found that knockdown of any individual DNMT only led to minor CD38 increase (**Fig. 1B**).

Given these findings, we therefore shifted our focus to genes that, when knocked down, led to CD38 decrease (left side of volcano plot in **Fig. 1B**). We reasoned this approach could still reveal important biological inputs that regulate the surface expression of CD38. Examining specific genes, we found that the transcription factor *SPI1* was the strongest hit besides *CD38* that, when knocked down, repressed surface CD38 expression. We also noted that *NFKB1* and *NFKB2* knockdown appeared to drive CD38 decrease. This finding was intriguing given the known importance of NF-κB signaling in myeloma proliferation and survival^37^. KEGG and GO analysis of genes whose knockdown significantly decreased CD38 showed enrichment for MAP kinase pathway and protein phosphorylation more broadly (**Supp. Fig. 1C**), suggesting key roles for intracellular signaling in regulating surface CD38.

Validation experiments with individual sgRNAs confirmed that *SPI1* knockdown strongly decreased CD38 surface expression by flow cytometry, with a lesser decrease in surface CD38 with *NFKB2* knockdown (**Fig. 1D**, **Fig. 2C, Supp. Fig. 2C-D**). These alterations also led to functional impacts. RPMI-8226 cells with knockdown of these genes showed significantly decreased NK cell lysis in ADCC assays (**Fig. 2D**). We further probed this dynamic *in vivo*, finding that RPMI-8226 cells with *SPI1* knockdown were relatively resistant to daratumumab in a murine model (**Fig. 2E**). We note that we attempted to expand these results to additional cell lines. However, our four other myeloma cell lines harboring the CRISPRi machinery^21^ all express extremely low levels of *SPI1* (**Supp. Fig. 2B**) and attempted knockdown in two of them (AMO1, KMS12PE) did not elicit any phenotype (not shown). Therefore, this finding suggests that SPI1 may play an important role in regulating CD38 expression in some myeloma tumors, but it is less likely to be a universal regulator.

### Epigenetic analysis suggests XBP1 as a key determinant of CD38 in primary myeloma tumors

Our CRISPRi results suggest that epigenetic and/or transcriptional regulation is a critical driver of surface CD38 levels. However, we do not know whether these specific hits in a myeloma cell line will extend to primary myeloma tumors. We therefore took a complementary approach to find potential transcriptional regulators of *CD38*. Using ATAC-seq data from 24 primary myeloma tumor samples^24^, we extracted open chromatin motifs near the *CD38* promoter (**Supp. Fig. 3A**) to identify a list of 46 transcription factors with potential binding sites at this locus (**Supp. Table 2**). We then correlated expression (via Pearson *R*) of these transcription factors with *CD38* expression across 664 primary patient tumors at diagnosis in the Multiple Myeloma Research Foundation CoMMpass database (research.themmrf.org; release IA13). In this analysis, we found the transcription factor most negatively correlated with *CD38* expression was *RARA* (**Fig. 3A**), consistent with our CRISPRi screen data, and underscoring the promise of ATRA as a co-treatment to increase *CD38*. Intriguingly, the transcription factor with strongest positive correlation was *XBP1* (**Fig. 3A-B**), a central driver of plasma cell identity^38^. *XBP1* also showed strong positive correlations with *CD38* in two other patient tumor gene expression datasets^39,40^ (**Supp. Table 2**).

**Figure 3.**
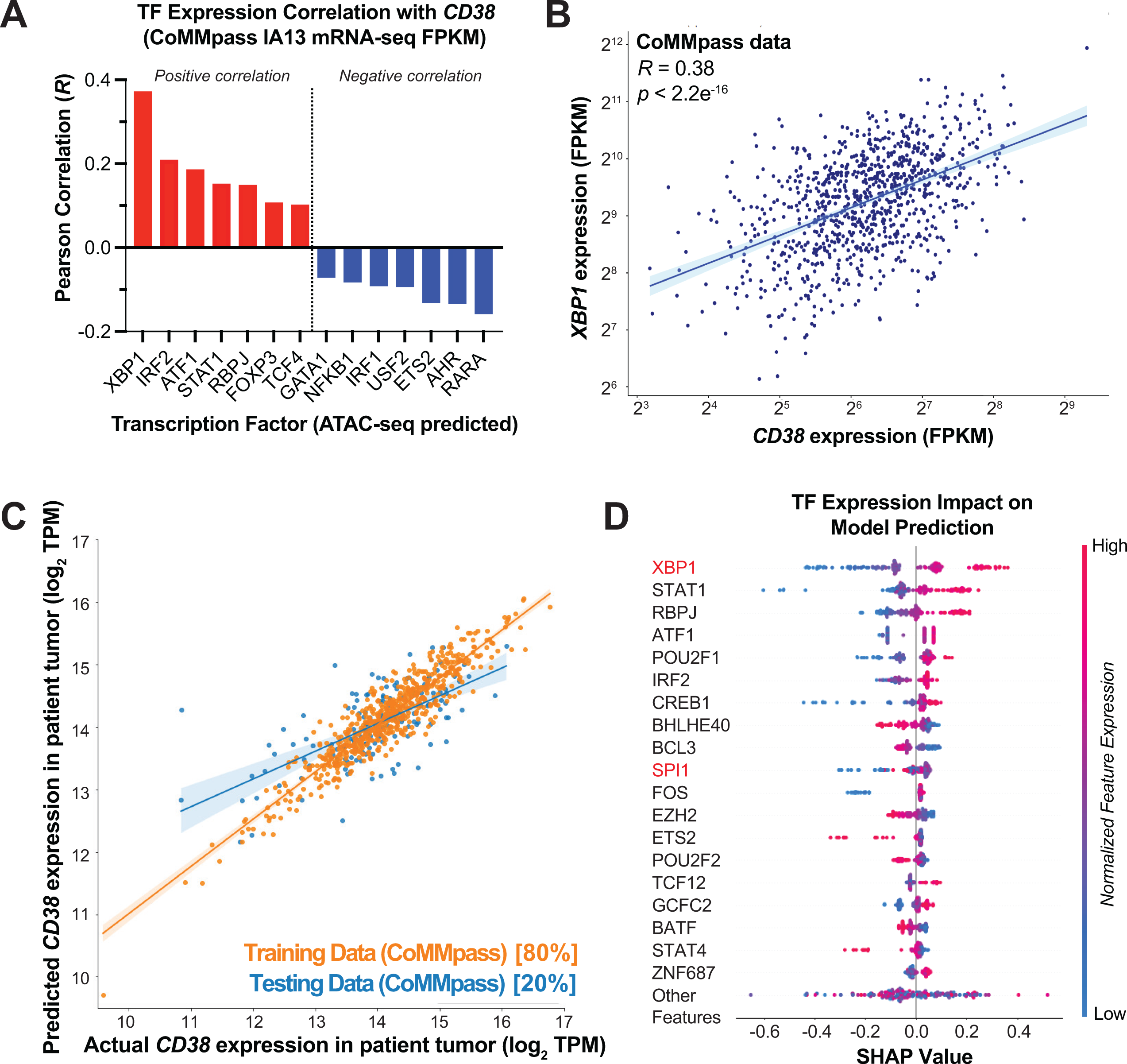
Patient-centered epigenetic analysis and machine learning predicts most potent transcriptional regulators of *CD38*. **A.** 46 transcription factors predicted to bind to the CD38 locus were derived from motif analysis of published ATAC-seq data (see Supplementary Figure 3). Gene expression of each transcription factor (TF) was correlated with *CD38* expression in the Multiple Myeloma Research Foundation (MMRF) CoMMpass database (release IA13), with RNA-seq data from CD138+ enriched tumor cells at diagnosis (*n* = 664 patients). Top predicted positive and negative regulators are shown based on Pearson correlation (*R*). **B.** CoMMpass RNA-seq data illustrates strong positive correlation between *XBP1* and *CD38* expression. **C.** XGBoost machine learning model was used to extract features of transcription factor gene expression that best-model CD38 expression in CoMMpass tumors (shown in log2 TPM (Transcripts per Million)). 80% of data was used as a test set with 20% left out as a training set. Coefficient of variation (*R*^2^) for predictive model = 0.49 after five-fold cross validation. **D.** Shapley Additive Explanations (SHAP) analysis indicates transcription factors whose expression most strongly impacts CD38 expression levels in CoMMpass tumors.

To further extend this analysis, we sought to build a predictive model which could estimate *CD38* transcript level as a function of transcription factor expression. We used an XGBoost method applied to CoMMpass mRNA-seq data to find weights of transcription factor expression that most-influence *CD38* levels in patient tumors. We first tested this analysis on 80% of patient data as a training set with 20% left out as test set. We found this model, solely based on transcription factor expression, could predict about half of the variance (coefficient of variation *R*^2^ = 0.49 using 5-fold cross validation) in test set *CD38* levels (**Fig. 3C**). Using model weights and Shapley Additive Explanations (SHAP) analysis (see Supplementary Methods) to determine transcription factors that have the greatest impact, either positive or negative, on CD38 expression, we found that *XBP1* played the strongest role overall. Other strong hits from both of our analyses included *IRF2*, *ATF1*, and *STAT1*. *SPI1* also appeared in the top 10 most relevant transcriptional regulators (**Fig. 3D; Supp. Fig. 3B**), consistent with our CRISPRi results, suggesting that SPI1 may play a key role in regulating *CD38* in a subset of tumors.

We further evaluated XBP1 given its prominent role in these two complementary bioinformatic analyses. In a prior dataset of shRNA knockdown of *XBP1* in myeloma plasma cells^41^, *CD38* mRNA was decreased ∼3-fold after *XBP1* silencing (**Supp. Fig. 3C**). This finding was consistent with results in our CRISPRi screen, where *XBP1* knockdown led to significant surface CD38 decrease (**Fig. 1B**). We further validated this relationship by using a doxycycline-inducible sgRNA construct for CRISPRi vs. *XBP1*, finding a clear dose-response between degree of XBP1 knockdown and loss of surface CD38 by flow cytometry (**Supp. Fig. 3D-G**). Supporting relevance of this link, a recent myeloma tumor whole genome sequencing study found that deletion of *XBP1* was one of the strongest determinants of clinical response to daratumumab^42^. We attempted to perform promoter activation assays to directly link XBP1 binding to *CD38* expression, but were unable to successfully generate a reporter construct that reflected all 8 loci where XBP1 may bind at *CD38* regulatory elements (**Supp. Fig. 3H**). Taken together, these results nominate XBP1 as a particularly strong determinant of surface CD38 in myeloma plasma cells, though future investigation will be required to validate a direct or indirect relationship to *CD38* transcription.

### No consistent large-scale remodeling of the myeloma surface proteome after CD38 downregulation

We next evaluated CD38 surface regulation from the perspective of monoclonal antibody therapy. In clinical samples, CD38 loss after daratumumab was accompanied by increases in CD55 and CD59, which may inhibit complement-dependent cytotoxicity and contribute to daratumumab resistance^5^. In preclinical studies, macrophage trogocytosis has been proposed as a mechanism contributing to CD38 loss after mAb treatment, that also leads to alterations in other surface antigens including CD138/SDC1 (ref.^7^). However, we hypothesized that given its enzymatic activity and role as a cellular differentiation marker^43^, loss of CD38 on its own may influence surface expression of other myeloma antigens. Such alterations may reveal new biology or (immuno)therapeutic vulnerabilities of CD38 mAb-treated disease.

To test this hypothesis, we employed a method we recently developed termed “antigen escape profiling”^27^. We used CRISPRi to transcriptionally repress *CD38* in RPMI-8226, AMO-1, and KMS12-PE myeloma cell lines, using this genetic approach to partially mimic loss of surface antigen seen after mAb therapy (**Fig. 4A**). We then performed Cell Surface Capture proteomics^26,27^ to uncover surface proteome alterations in a relatively unbiased fashion. Across cell lines, analyzed in biological triplicate with *CD38* knockdown vs. non-targeting sgRNA, we quantified 897 proteins annotated as membrane-spanning in Uniprot (minimum of two peptides per protein) (**Supp. Table 3**). As a positive control, in all lines we found that the strongest signature was decrease of CD38 itself (**Supp. Fig. 4A-C**). However, when aggregating proteomic data, we found no significant alterations in any surface antigens beyond CD38 itself (**Fig. 4B**). Integration with RNA-seq data revealed only THY-1/CD90 as upregulated >3-fold at both the mRNA and surface proteomic level after *CD38* knockdown (**Fig. 4C**). Intriguingly, CD90 is known as a marker of “stemness” in early hematopoietic lineage cells that is lost when CD38 expression is increased^44^. CoMMpass analysis also confirmed increased *THY1* expression in tumors with lower *CD38* (**Supp. Fig. 4D**). However, further validation as to whether CD90 is truly altered after CD38 mAb will require pre- and post-treatment clinical specimens, beyond the scope of our work here. Overall, we conclude that loss of CD38 in isolation leads to limited remodeling of the myeloma surface proteome.

**Figure 4.**
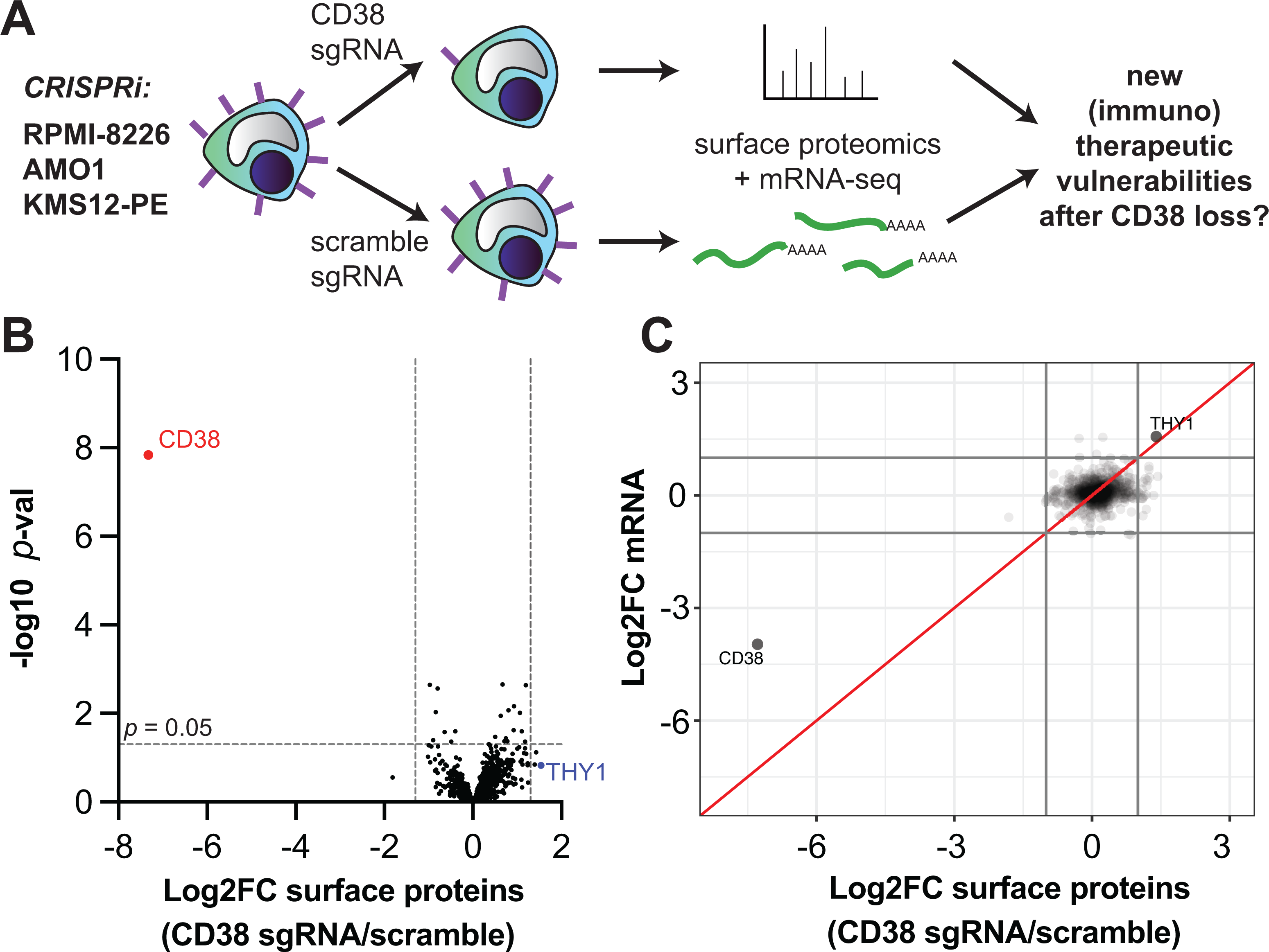
Minimal alterations of the myeloma cell surface proteome after CD38 loss. **A.** Schematic of “antigen escape profiling” approach to reveal new cell-surface therapeutic vulnerabilities in the context of CD38 downregulation. **B.** Cell surface capture proteomics comparing *CD38* knockdown vs. non-targeting sgRNA control, with aggregated data across three cell lines (CRISPRi-expressing RPMI-8226, AMO1, and KMS12-PE; *n* = 3 replicates per cell line per sgRNA) reveals minimal changes in the cell surface proteome beyond CD38 knockdown at significance cutoff of p < 0.05 and log_2_ fold-change >|1.5|. **C.** Integrated analysis of cell surface proteomics and mRNA-seq (*n* = 2 per cell line per guide) across three cell lines reveals the only consistent change at both protein and transcript level after *CD38* knockdown is *THY1*/CD90 upregulation. Log_2_ fold-change cutoff = |1.5|.

### Integrated surface proteomic and transcriptional analysis suggests ATRA is highly specific in CD38 upregulation

Data from our group^16^ and others^12–15^ have suggested that several small molecules can increase myeloma surface CD38. However, the broader impacts of these agents on membrane antigens beyond CD38 have not been directly compared. We performed integrated cell surface proteomics and transcriptional analysis of RPMI-8226 cells treated with 10 nM all-trans retinoic acid (ATRA), 2 μM azaciditine (Aza), and 10 nM Panobinostat (Pano), all treated for 72 hr, in comparison to DMSO (**Supp. Table 4**). These doses are chosen as they have been previously published to significantly increase myeloma surface CD38 by flow cytometry^12,13,16^. In this integrated analysis we found much broader impacts of azacytidine and panobinostat than ATRA on the “surfaceome” of plasma cells, beyond increasing CD38 (**Fig. 5A**). These results suggest that, at doses driving CD38 upregulation, for Aza or pano altering CD38 is just a small component of their impact on myeloma tumor cells, whereas ATRA is much more specific in driving CD38 upregulation.

**Figure 5.**
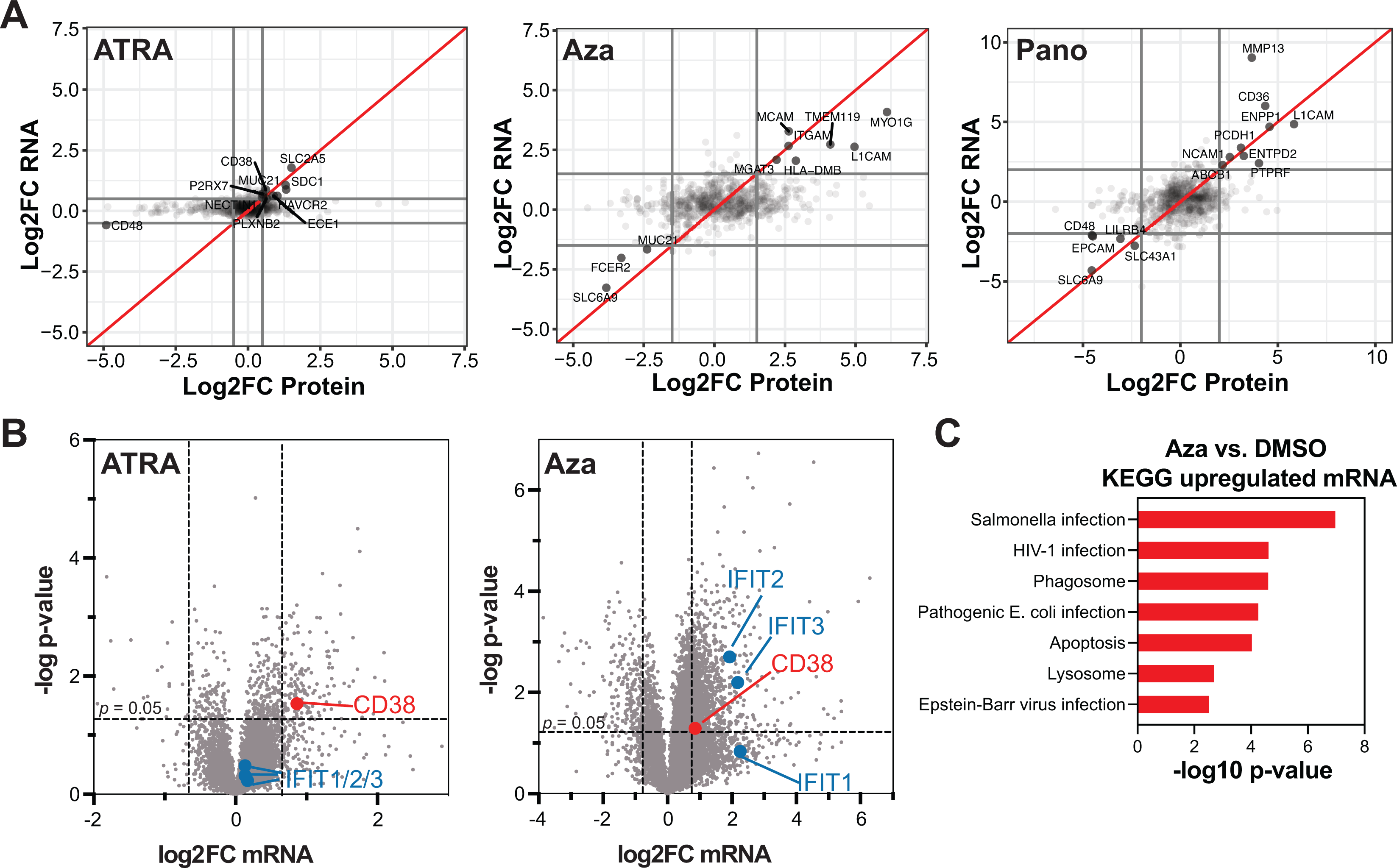
ATRA drives CD38 upregulation with limited additional cellular impact while Aza leads to a broad interferon-mediated response. **A.** Integrated mRNA-seq (*n* = 2 per drug treatment) and cell surface proteomics (*n* = 2 per drug treatment) across RPMI-8226 treatment with 10 nM all-trans retinoic acid (ATRA), 2 μM azacytidine (Aza), and 10 nM panobinostat (Pano). All plots are in comparison to control replicates treated with 0.1% DMSO. Doses chosen are based on those previously published to lead to CD38 upregulation for each agent. Data points shown are for proteins and genes corresponding to Uniprot-annotated membrane-spanning proteins. Log_2_-fold change cutoffs shown at |0.5| for ATRA and |2.0| for Aza and Pano to increase clarity of plots given many fewer changed genes with ATRA treatment. **B.** RNA-seq for same samples with ATRA or Aza treatment vs. DMSO but here showing all mapped genes, not just those annotated as membrane-spanning. Significance cutoff at *p* < 0.05 with log_2_ fold-change cutoff set at |0.8| to illustrate prominent differences above this level in transcriptome alteration after either ATRA or Aza treatment. **C.** KEGG analysis of genes from RNA-seq dataset meeting cutoff criteria of *p* < 0.05 and log_2_ fold-change >0.8 after Aza treatment.

Toward understanding how CD38 is modulated after drug treatment, in our previous work^16^ we noted that the mechanism of *CD38* increase after Aza treatment was unclear. We thus further investigated the global transcriptional response (i.e. not limited to membrane) after Aza (**Supp. Table 4**). Prior studies have suggested that Aza anti-tumor effect is largely mediated by reactivation of endogenous retroviruses stimulating a tumor-autonomous interferon response^45,46^. Consistent with this work, we found a pronounced increase in interferon-responsive genes after Aza, but not ATRA, including *IRF1*, *IFITM1*, *IFITM2*, and *IFITM3* (**Fig. 5B**). KEGG analysis also confirmed this effect (**Fig. 5C**). Given evidence across multiple systems that interferon upregulates *CD38* expression^47,48^, our transcriptional profiling thus also supports an interferon-based mechanism driving surface CD38 increase in plasma cells after Aza treatment.

### Plasma cell proliferative signaling pathways are inhibited by mAb binding to CD38

In our final set of experiments related to targeting surface CD38, we were intrigued as to whether binding of a therapeutic mAb leads to specific cellular phenotypes within myeloma plasma cells. For example, isatuximab is known to directly lead to apoptosis of plasma cells^49^, and daratumumab can do so after crosslinking^50^. However, the mechanism underlying this transduction of extracellular mAb binding to intracellular phenotype remains unclear. In addition, our CRISPRi screen data (**Fig. 1B and Supp. Fig. 1C**) suggests that surface CD38 expression may be strongly impacted by intracellular phospho-signaling pathways.

Therefore, we used unbiased phosphoproteomics by mass spectrometry to probe intracellular signaling effects driven by CD38 mAb binding. In RPMI-8226 cells we compared 20 μM daratumumab treatment vs. IgG1 isotype control. We chose a time point of 20 minutes of treatment given known rapid alterations in signaling pathways in similar phosphoproteomic experiments^51^. In total, across triplicate samples we quantified 5430 phosphopeptides (**Supp. Table 5; Supp. Fig. 5A**). Analyzing phosphopeptide changes by Kinase Substrate Enrichment Analysis (KSEA)^52^, we were intrigued to find downregulation of phosphorylation motifs consistent with both cyclin-dependent kinases as well as several kinases of the MAP kinase pathway (**Fig. 6A**). Downregulation of phosphorylation on several central nodes in the MAP kinase as well as AKT pathway was also apparent via KEGG analysis (**Supp. Fig. 5B**). Across a time course we further confirmed effects on MAP kinase pathway (reported by phosphorylation of MAPK (ERK1/2), a key node in this response) and AKT signaling after daratumumab treatment via Western blotting in RPMI-8226 and MM.1S cell lines, respectively (**Fig. 6B**). While the absolute value of changes in MAPK signaling are modest, both by phosphoproteomics and Western blot, these results indicate that daratumumab binding to CD38 can at least partially inhibit this central proliferative pathway within myeloma tumor cells, and thus may form a component of daratumumab’s anti-tumor effect.

**Figure 6.**
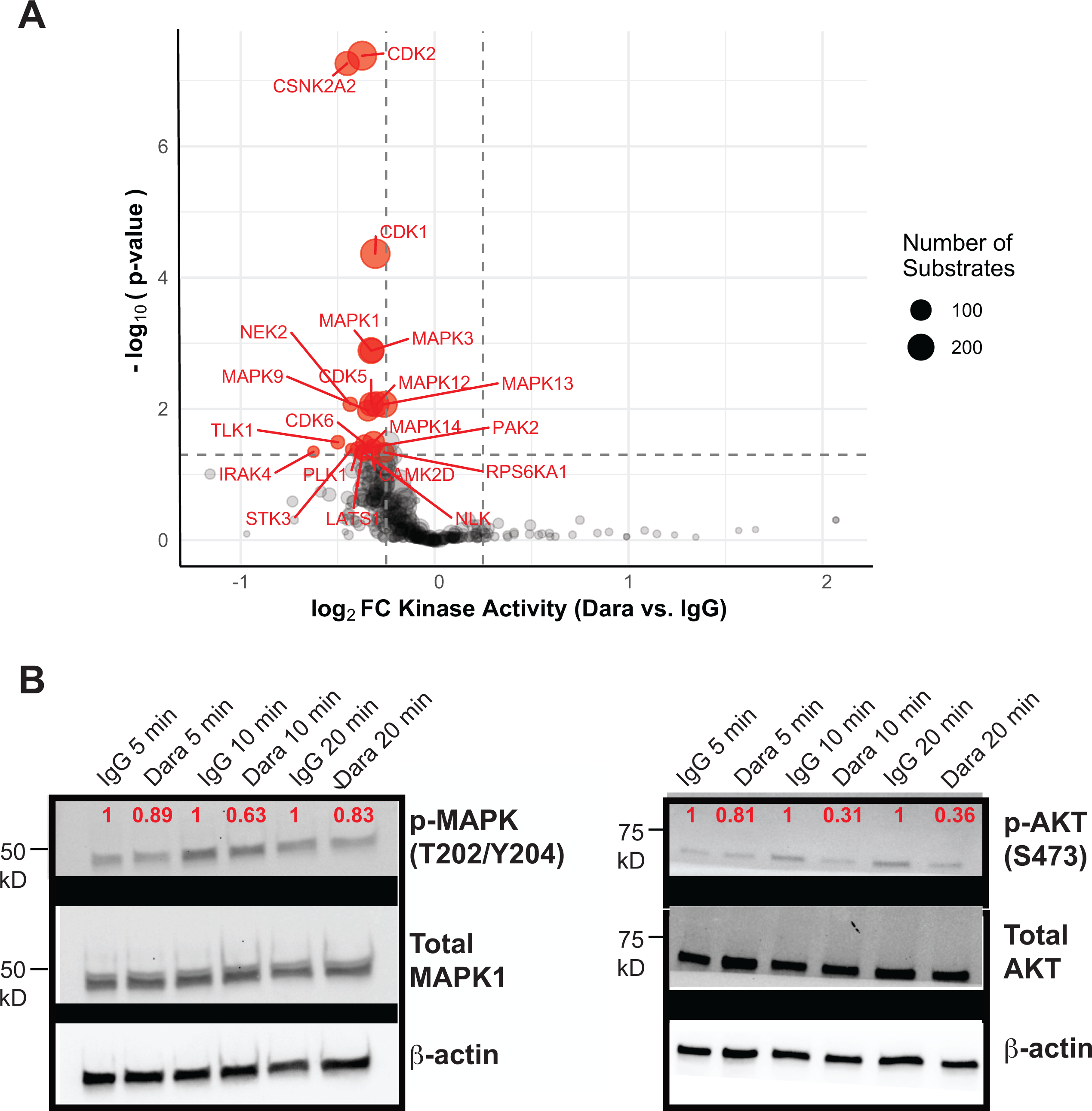
Unbiased phosphoproteomics reveals downregulation of proliferative signaling after daratumumab treatment. **A.** RPMI-8226 cells were treated with 20 μM daratumumab or IgG1 isotype control for 20 minutes (*n* = 3 each) and then harvested for unbiased phosphoproteomics with immobilized metal affinity chromatography enrichment for phosphopeptide enrichment. Plot displays results of Kinase Substrate Enrichment Analysis, indicating modest decrease in phosphorylation of numerous predicted substrates of MAPK pathway kinases as well as cyclin-dependent kinases (cutoffs of *p* < 0.05, log_2_ fold-change >|0.5|). **B.** Western blot in RPMI-8226 of MAPK (ERK1/2) (Thr202/Tyr204) relative to total MAPK demonstrates modest decrease in MAPK phosphorylation after 5, 10, or 15 min daratumumab (Dara) treatment; magnitude of change normalized to IgG1 control at each time point (red) appears consistent with phosphoproteomic data. **C.** Western blot of MM.1S cells treated with daratumumab and blotted for p-AKT (Ser473) and total AKT, with quantification of p-AKT relative to total AKT and normalized to IgG1 at each time point. All images representative of two independent Western blots.

## Discussion

Our studies here present a “multi-omics” view of therapeutically targeting CD38 in multiple myeloma. Our integrated functional genomics and epigenetic analysis point to the central role of transcriptional regulators in governing CD38 surface expression. Using surface proteomics, we further identify that loss of CD38 in isolation is unlikely to drive large changes in the “surfaceome”, while known pharmacologic strategies to increase CD38 have largely divergent impacts on other surface antigens. Finally, unbiased phosphoproteomics reveals that binding of anti-CD38 mAb can impair intracellular proliferative signaling within plasma cells.

Our CRISPRi screen illustrated the central role of numerous transcription factors, such as SPI1, HEXIM1, and TLE3, in regulating surface CD38. This functional genomic study suggests that regulation of surface CD38 largely occurs at the transcriptional, as opposed to protein trafficking, level. This finding was in sharp contrast to our prior CRISPRi results with BCMA, where we found that post-transcriptional mechanisms, such as proteolytic cleavage by γ-secretase and protein trafficking via the SEC61 translocon, played the strongest roles in determining surface BCMA levels^21^.

Another recent study used genome-wide CRISPR deletion screening to find genes that, when knocked out, could abrogate IL-6-mediated downregulation of surface CD38 (ref.^15^). The strongest hits in this prior study included the transcription factors *STAT1* and *STAT3*, demonstrating a role for JAK-STAT signaling in regulating tumor CD38 expression within the bone marrow microenvironment^15^. In support of this notion, our integrated epigenetic and machine learning analyses, extracted from bone marrow-derived patient tumor samples, also support a critical role for STAT1 in governing surface CD38. However, in our CRISPRi screen in an *in vitro* monoculture system, neither *STAT1* or *STAT3* affected CD38 surface expression (**Fig. 1B**). This result suggests that JAK-STAT signaling may not play a major role in CD38 regulation in the absence of exogenous tumor stimulation. This finding illustrates the complementary nature of our genome-wide screen to that previously published under the context of IL-6 stimulation^15^.

Toward the goal of finding key regulators of CD38 that were not previously known, our epigenetic and machine learning approaches suggest that *XBP1* is a critical regulator of plasma cell *CD38*. To our knowledge, there are not currently any known pharmacologic mechanisms to potentiate XBP1 activity. Given the important role of *XBP1* splicing in myeloma plasma cells^53^, future work will investigate the role spliced vs. unspliced XBP1 in specifically regulating CD38, as this strategy may provide new avenues for CD38 manipulation. Future work will also investigate the role of *XBP1* deletion in determining clinical response to daratumumab^42^.

Given that plasma cells demonstrate frequent loss of CD38 after daratumumab treatment^5^, a pressing question is whether CD38-low, daratumumab-resistant cells have novel immunotherapeutic vulnerabilities. However, our recently-described strategy of “antigen escape profiling”^27^ – CRISPRi knockdown followed by unbiased cell surface proteomics (**Fig. 4**) – suggests that other surface antigens on plasma cells do not exhibit consistent changes due to CD38 downregulation alone. This finding supports the notion that alterations in surface proteins found after mAb treatment on patent tumors, such as increases in CD55 and CD59 (ref.^5^), are caused by other therapy-induced selective pressure within the tumor microenvironment, not CD38 loss.

While *in vitro* assays have suggested a strong relationship between CD38 antigen density and either NK-cell^54^ or macrophage-mediated antibody-dependent cell killing^6^, these experiments cannot readily take into account the critical role of the immune microenvironment in determining daratumamab response or resistance^8^. Furthermore, even with the higher specificity of ATRA, there is the potential to alter CD38 expression on other hematopoietic cells, which may impact clinical responses to daratumumab^17^. Notably, current clinical data is most consistent with pre-treatment tumor CD38 antigen density positively correlating with daratumumab depth of response^5,11^. Analysis of transcriptional data in CoMMpass demonstrates that while increased *CD38* expression was associated with improved outcomes in all patients, only those treated with daratumumab showed a divergence in survival curves coinciding with the median time of initial daratumumab treatment (**Supp. Fig. 6**). Pharmacologic manipulation of CD38 density on tumor cells may ultimately be most fruitful pre-treatment rather than in the context of daratumumab resistance. Similar strategies may also be most beneficial for other CD38-targeting immunotherapeutics^2^.

Also directly related to mAb therapeutic effects, our unbiased phosphoproteomic results suggest that daratumumab binding to CD38 can directly decrease signaling along the MAP kinase and PI3K-AKT pathways. It remains to be investigated whether this inhibition of central proliferative signaling pathways plays a role in the anti-tumor effect of daratumumab in patients.

In terms of limitations of our work, the most prominent is that the many of our studies are derived from large-scale “omics” experiments in myeloma cell lines. There may be biological differences between our findings *in vitro* and primary tumors growing within the bone marrow microenvironment.

Taken together, our multi-omic studies comprise a resource that reveals new insight into the genetic, epigenetic, and pharmacologic regulation of surface CD38 in myeloma plasma cells. We anticipate these findings will have utility in deriving new strategies to enhance CD38-targeting therapies in myeloma, including mAbs in current clinical practice as well as emerging antibody and cellular therapies^2^. The technologies described here also comprise a blueprint to comprehensively assess determinants of surface antigen regulation, and impacts of associated therapeutic manipulation, that could be applied across targets in hematologic malignancies.

## Acknowledgements

We thank Ruilin Tian for assistance in modifying MAGeCK scripts for the analysis here. This work was supported by grants K08 CA184116, R01 CA226851, and the UCSF Stephen and Nancy Grand Multiple Myeloma Translational Initiative (to A.P.W.), NCI P30 CA082103 supporting the Preclinical Therapeutics Core facility (managed by V. Steri and B.H.), and K99/R00 CA181494 and a Stand Up to Cancer Innovative Research Grant (to M.K.).

## Author Contributions

P.C., P.R., M.K., and A.P.W. conceived and designed the study. P.C., C.K., B.P.E., A. Kang, A. Kishishita, S.R., J.C.P., O.G., L.S., Y-H.T.L., P.R., N.P., and M.M. performed experiments. P.C., B.P.E., N.P., Y-H.T.L., B.J.B., and P.R. analyzed data. D.W., P.P., V. Steri, and B.H. performed murine studies. H.G. analyzed patient epigenetic data. V. Sarin performed machine learning analysis. P.C. and A.P.W. wrote the manuscript with input from all authors.

