## Supplementary Materials for "Functional multi-omics reveals genetic and pharmacologic regulation of surface CD38 in multiple myeloma"

Choudhry et al.

**Supplementary Figures 1-6**

**Supplementary Methods**

**Supplementary Table Legends**

**(Supplementary Tables attached as separate Excel spreadsheets)**

**Supplementary References**

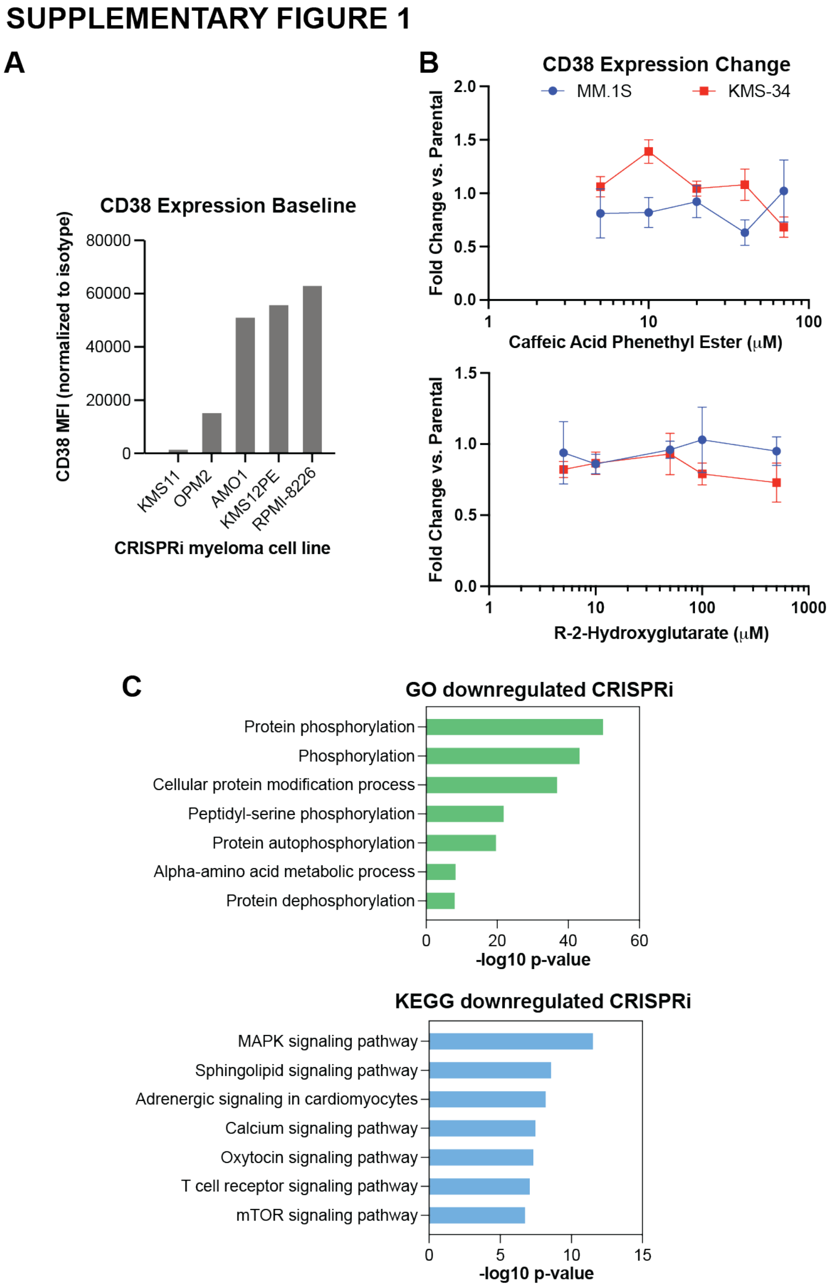


**Supplementary Figure 1. Additional results of CRISPRi screen. A.** Validation that RPMI-8226 cells express robust levels of CD38 in comparison to other myeloma cell lines harboring the CRISPRi machinery. *n* = 1 replicate. **B.** Treatment of MM.1S or KMS11 cells with caffeic acid phenethyl ester (CAPE) or R-2-Hydroxyglutarate have no consistent impact on surface CD38 fold-change measured by flow cytometry. *n* = 3 technical replicates. Mean +/- S.E.M. displayed. **C.** GO and KEGG analysis of genes that when knocked down significantly decrease surface CD38, from data in **Fig. 1B**.

### SUPPLEMENTARY FIGURE 2

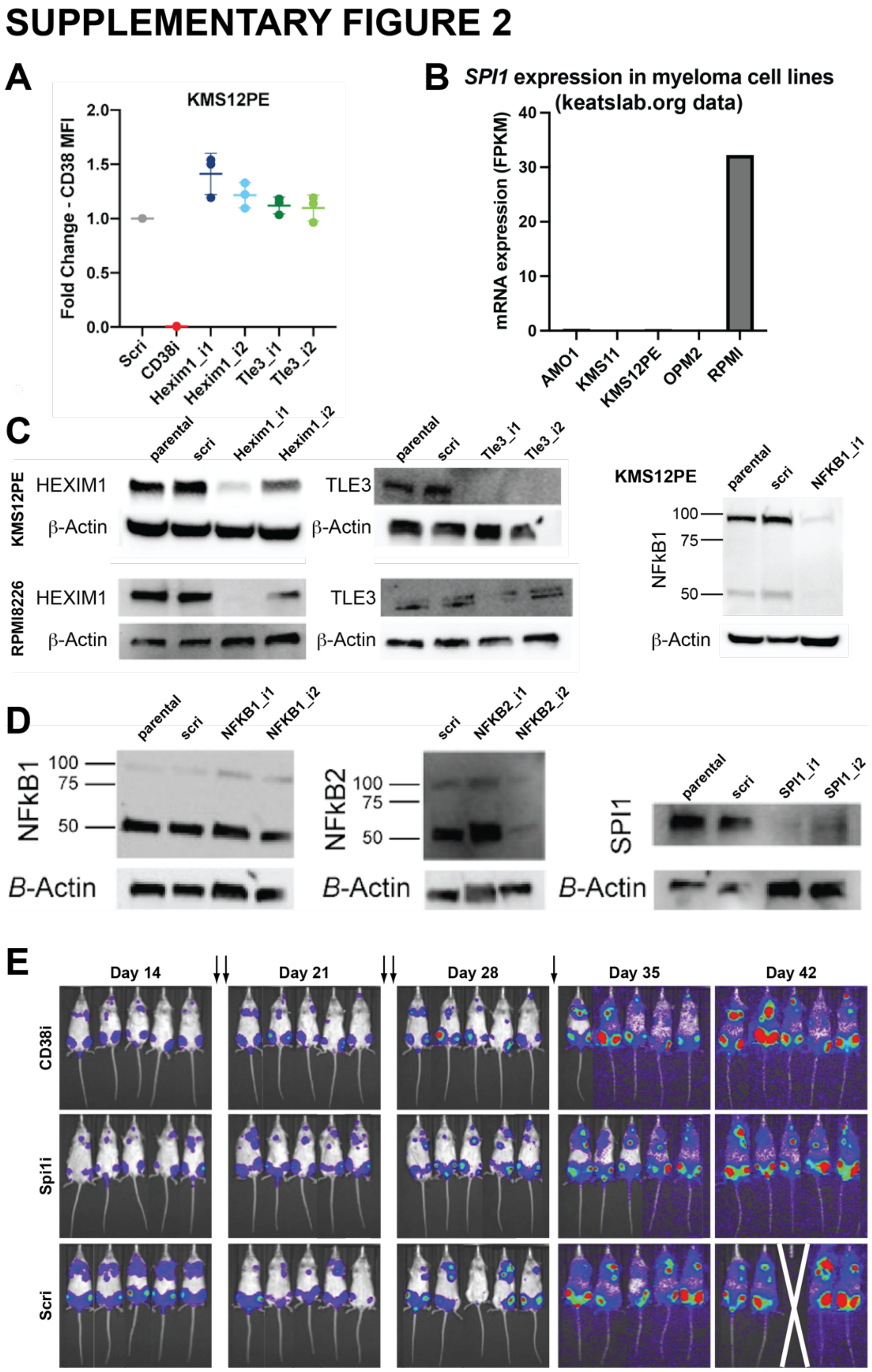


**Supplementary Figure 2. Supporting data for knockdown and validation experiments from CRISPRi screen. A.** Knockdown of HEXIM1 and TLE3 in KMS12-PE cells also leads to trend of increased surface CD38 measured by flow cytometry, potentially consistent with CRISPRi screen in RPMI-8226. Scri = non-targeting control sgRNA. *n* = 3 technical replicates. Mean +/- S.E.M. displayed. No significant differences were observed by one-way ANOVA. **B.** Publicly available mRNA-seq data (keatslab.org) indicates that of available cell lines harboring CRISPRi machinery, only RPMI-8226 expresses *SPI1*, precluding validation of *SPI1* knockdown in additional cell lines. **C.** Western blots demonstrating relative efficacy of different sgRNA's targeting HEXIM1 and TLE3 across KMS12PE and RPMI-8226 lines, and NFkB1 knockdown in KMS12PE. Parental = parental CRISPRi cell line with no sgRNA transduced. **D.** Western blots demonstrating relative CRISPRi knockdown of NFKB1, NFKB2, and SPI1 in RPMI-8226 cells. **E.** Representative bioluminescence imaging of NSG mice used in murine studies. Note that both *CD38* knockdown and *SPI1* knockdown impair initial tumor growth *in vivo* (decreased signal at Day 14, prior to daratumumab treatment). *Right* shows fold-increase in bioluminescence signal at each time point for each sgRNA compared to Day 14 signal. Arrows indicate timing of daratumumab dosing (200 ug/dose I.P.). *n* = 5 animals per arm.

### SUPPLEMENTARY FIGURE 3

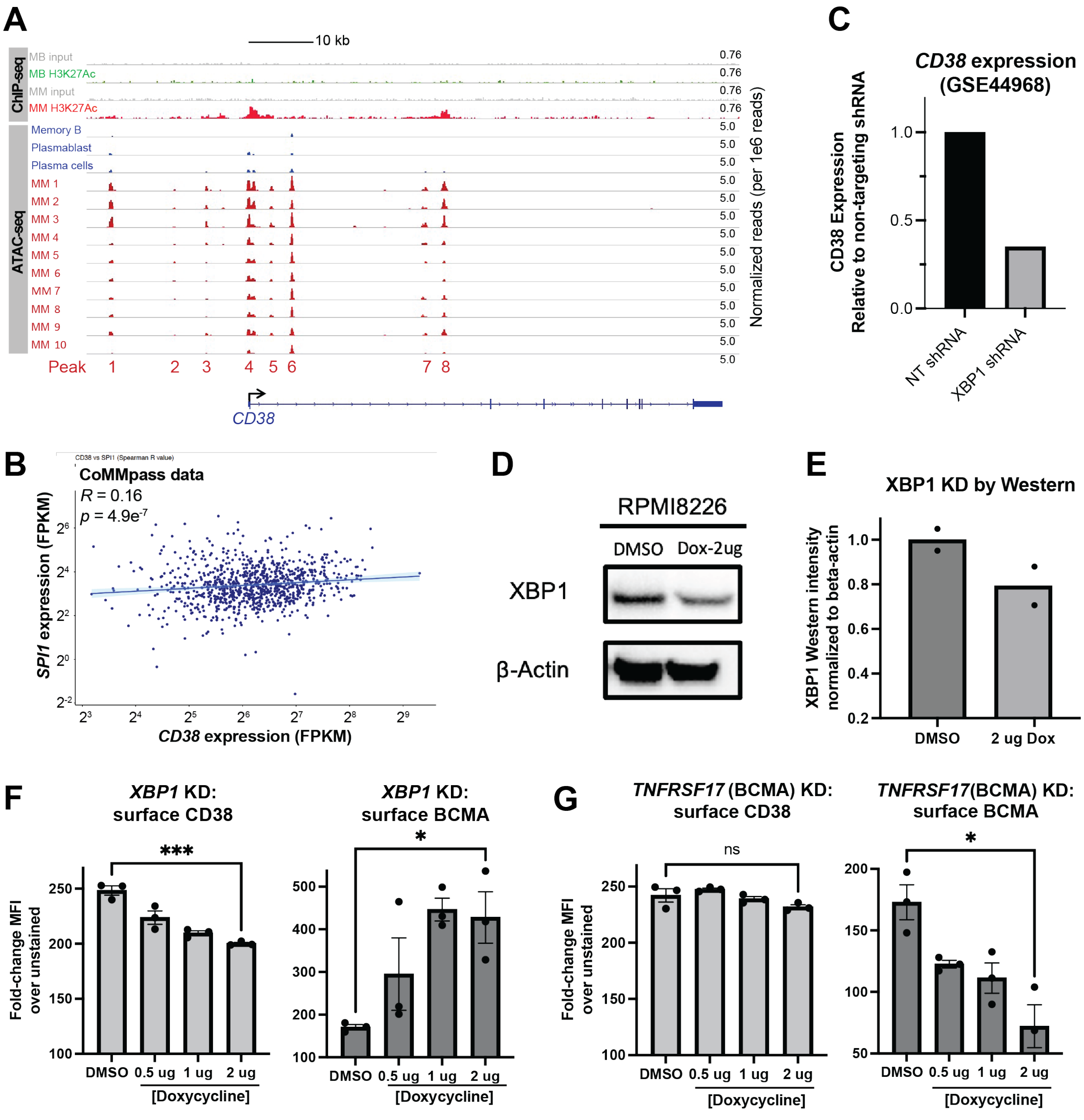


H

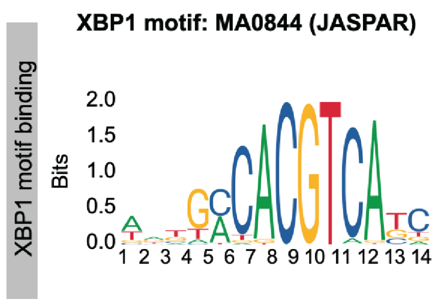


| Motif ID | Gene | Chr. | Strand | Start | End | p-value | Matched Sequence | ATAC peak |
|---|---|---|---|---|---|---|---|---|
| MA0844.1 | XBP1 | chr4 | - | 15758195 | 15758208 | 0.00046 | TGCAACACGTCCTA | peak.1 |
| MA0844.1 | XBP1 | chr4 | - | 15768013 | 15768026 | 0.00115 | TCAGCCACATCCTC | peak.2 |
| MA0844.1 | XBP1 | chr4 | + | 15767969 | 15767982 | 0.00182 | GCTGAAAAGTCACC | peak.2 |
| MA0844.1 | XBP1 | chr4 | - | 15773087 | 15773100 | 0.00089 | TCTGCTACATCACT | peak.3 |
| MA0844.1 | XBP1 | chr4 | + | 15780134 | 15780147 | 0.00101 | TGATCCTCGTCGTG | peak.4 |
| MA0844.1 | XBP1 | chr4 | + | 15780230 | 15780243 | 0.00176 | CGCGATGCGTCAAG | peak.4 |
| MA0844.1 | XBP1 | chr4 | + | 15780735 | 15780748 | 0.00201 | GAGGACACTTAAGT | peak.4 |
| MA0844.1 | XBP1 | chr4 | + | 15783319 | 15783332 | 0.00227 | ACCTTCACATCACA | peak.5 |
| MA0844.1 | XBP1 | chr4 | - | 15786456 | 15786469 | 0.00016 | CTGACCTCGTCATC | peak.6 |
| MA0844.1 | XBP1 | chr4 | + | 15807627 | 15807640 | 0.00113 | AGGGACACATTATG | peak.7 |
| MA0844.1 | XBP1 | chr4 | - | 15810559 | 15810572 | 0.00064 | CTAGACACTTCACC | peak.8 |
| MA0844.1 | XBP1 | chr4 | - | 15811316 | 15811329 | 0.00234 | AGTGTCACGGCAAT | peak.8 |

**Supplementary Figure 3. Supporting data for epigenetic analysis of transcription factors regulating *CD38* expression. A.** Called peaks from ChIP-seq and ATAC-seq (data from ref.[1] at the *CD38* locus, across normal memory B-cells (MB), plasmablasts, normal plasma cells, and malignant plasma cells derived from multiple myeloma (MM) patients (ten illustrated here; 24 total included in study). ATAC-seq peak numbers correspond to those noted in **Supp. Table 2**. **B.** Pearson correlation of CoMMpass tumor RNA-seq gene expression for *SPI1* and *CD38*. *p*-value by associated two-tailed *t*-test. **C.** Microarray gene expression data from ref.[2] demonstrates markedly decreased *CD38* expression after shRNA knockdown of *XBP1* in myeloma plasma cells. **D.** XBP1 Western blot confirms knockdown of *XBP1* via doxycycline-inducible sgRNA and CRISPRi, after 2 ug doxycycline for 72 hrs. Representative image of $n = 2$ biological replicates. **E.** Quantification of XBP1 Western blot results after inducible knockdown, normalized to beta-actin. Note that statistical analysis cannot be performed given less than three replicates. **F.** *Left:* Flow cytometry for CD38 after induced *XBP1* knockdown for 72 hrs at the doxycycline doses noted; 0.1% DMSO used as negative control. $n = 3$ biological replicates. *Right:* Same design as (F) but with flow cytometry for BCMA after XBP1 knockdown. **G.** *Left:* Same design as (F) but with flow cytometry for CD38 after induced *TNFSFR17* (encoding BCMA) knockdown, as a negative control. *Right:* Same design as (F) but with BCMA flow cytometry after *TNFSFR17* knockdown, as a positive control. Mean +/- S.E.M. displayed. *p*-values by two-tailed *t*-test. ***<0.005; **<0.01; *<0.05. **G.** MEME FIMO motif search tool[3] was used for scanning occurrences of XBP1 motif in the ATAC peak regions in the *CD38* gene. Sequences which matches XBP1 motif with p<0.0025 was shown. XBP1 motif logo and frequency matrix was obtained from JASPAR (https://jaspar.elixir.no/matrix/MA0844.1/).

### SUPPLEMENTARY FIGURE 4

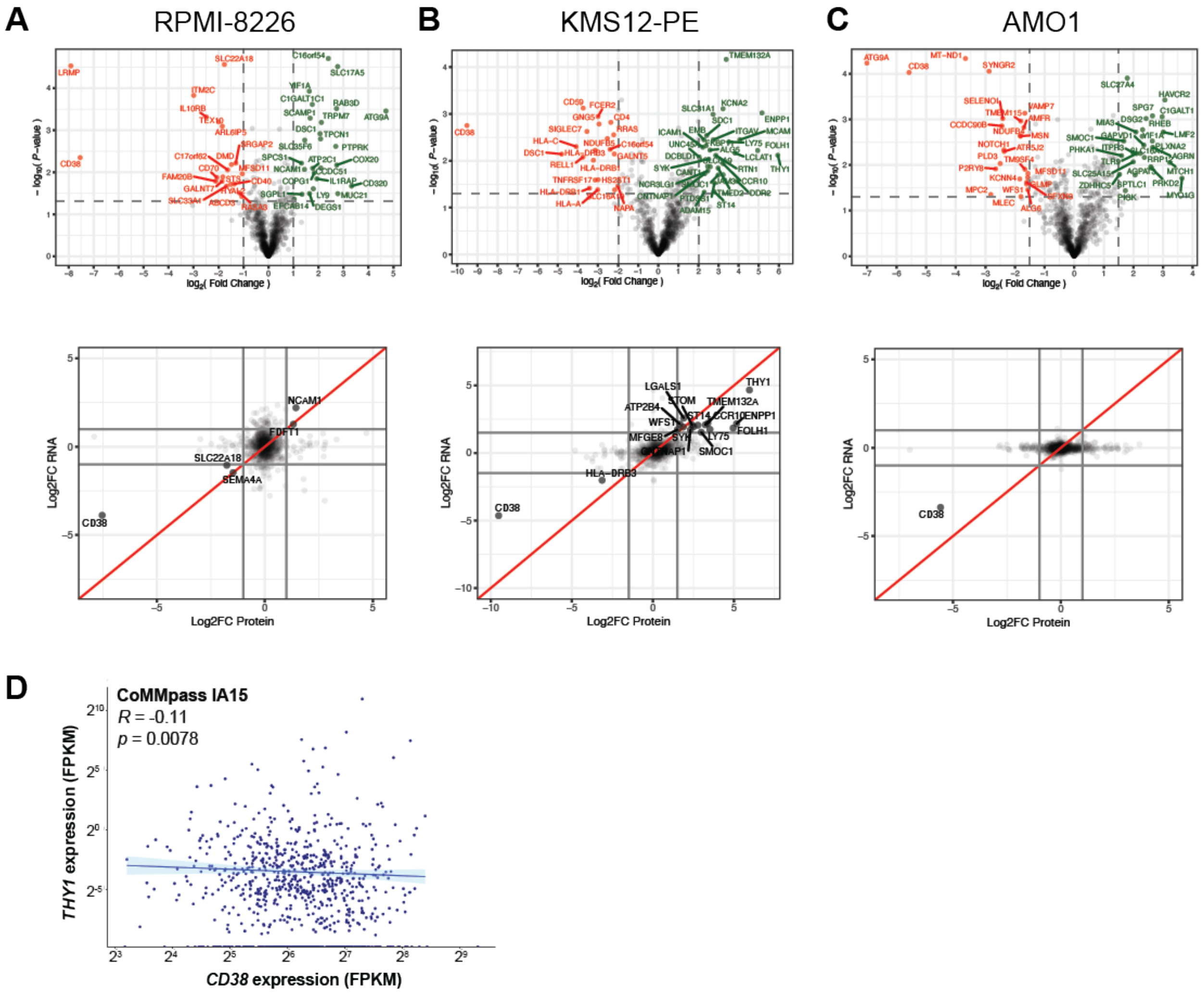


**Supplementary Figure 4. Supporting data for individual cell lines examined by "Antigen Escape Profiling". A.-C.** Surface proteomics (*top*) and integrated RNA-seq and surface proteomics (*bottom*) for each of the cell lines (RPMI-8226, KMS12-PE, AMO1) included in the aggregated analysis of **Fig. 4**. Each plot shows comparison of *CD38* sgRNA knockdown vs. non-targeting sgRNA control. $n = 3$ for proteomics, $n = 2$ for RNA-seq. For volcano plots significance cutoffs are $\log_2$-fold change >|1| and $p < 0.05$ by two-sided *t*-test. **D.** Pearson correlation of CoMMpass tumor RNA-seq gene expression for *THY1* and *CD38*. *p*-value by associated two-tailed *t*-test.

### SUPPLEMENTARY FIGURE 5

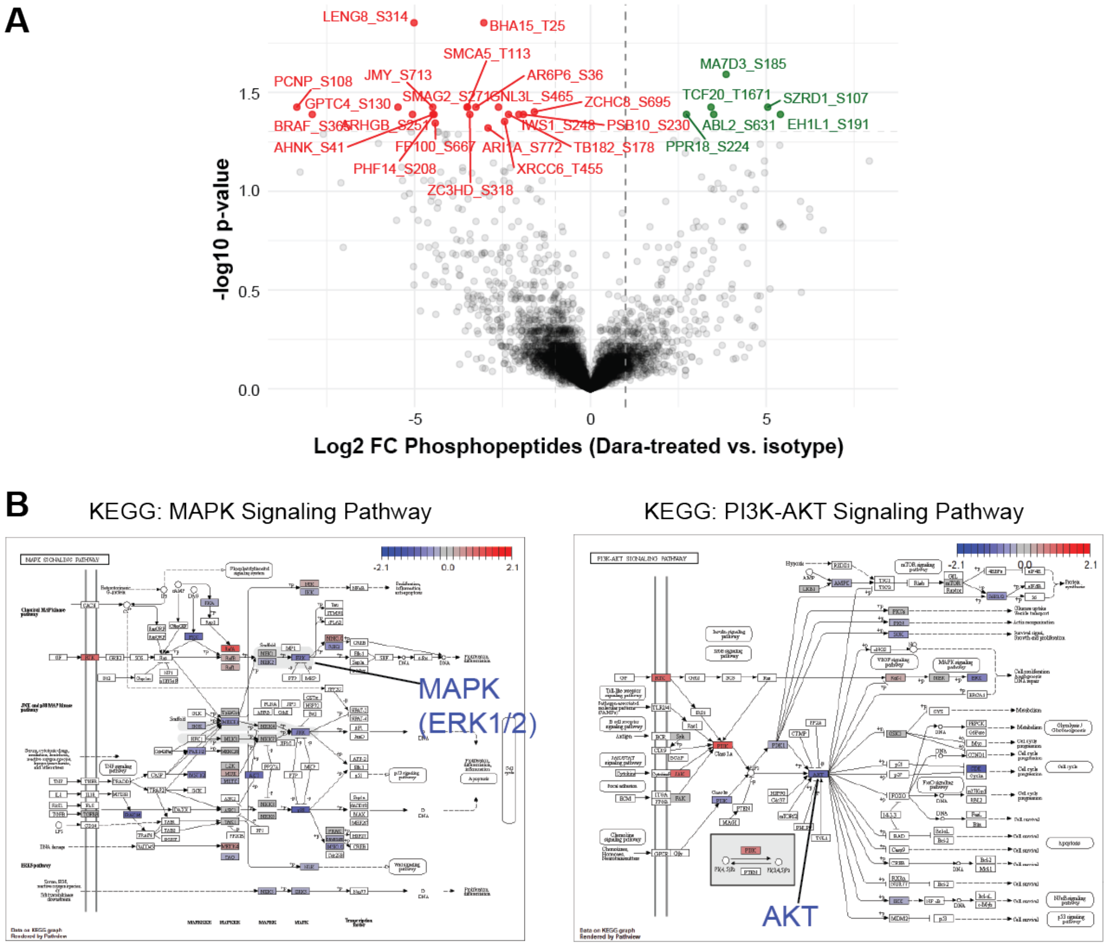


**Supplementary Figure 5. Additional analysis of phosphoproteomics experiments after daratumumab treatment. A.** Volcano plot of fold-change of individual phosphopeptides detected in RPMI-8226 after 20 min of 20 uM daratumumab treatment, relative to IgG1 isotype control. Displayed significance cutoffs of $p < 0.05$ by two-tailed $t$-test and $\log_2$ fold-change >|1|. **B.** KEGG pathways displaying alterations in phosphopeptides detected for included proteins across the MAPK and PI3K-AKT pathways. Red coloration indicates increased phosphorylation; blue indicates decreased. Phosphoproteomics shows decreased phosphorylation of MAPK1/ERK and AKT, consistent with Western blot results in **Fig. 6**.

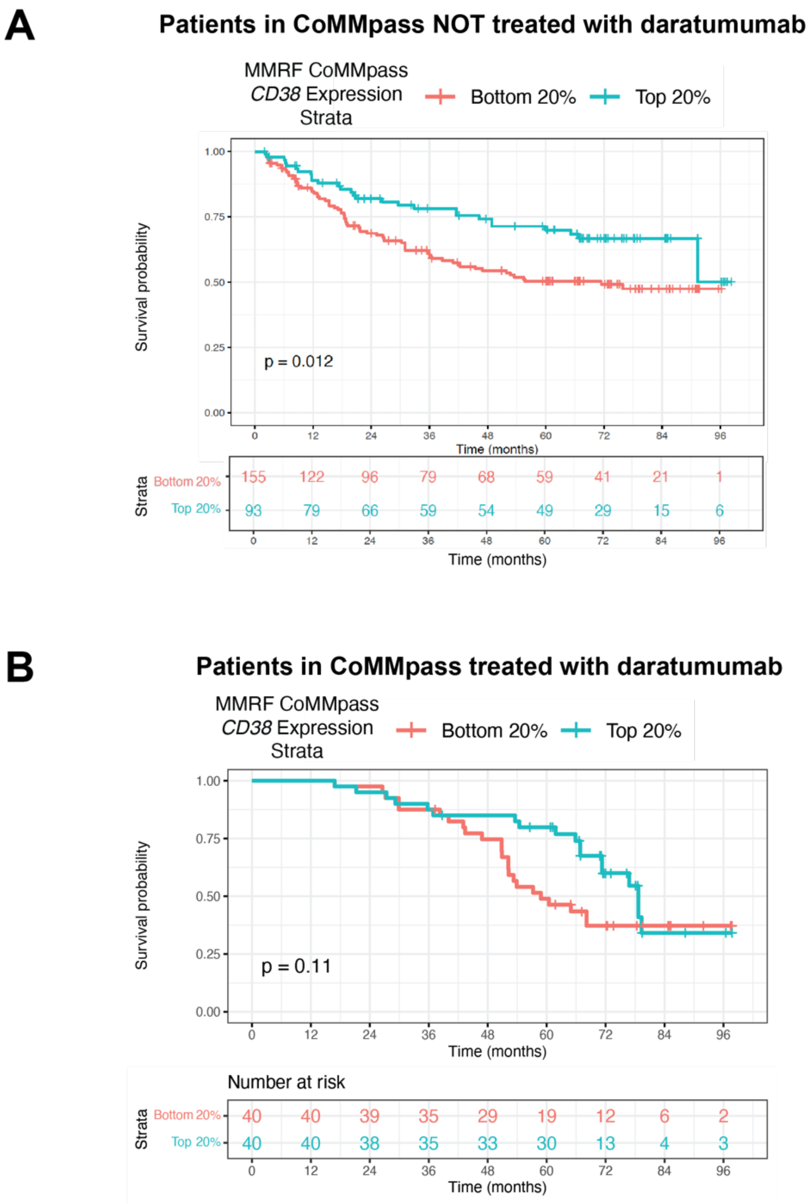


**Supplementary Figure 6. Overall survival in CoMMpass for patients recorded to receive daratumumab or not receive daratumumab, as a function of tumor *CD38* expression. A.** Correlation between tumor *CD38* expression quintile and overall survival for CoMMpass patients (release IA19) who did not receive daratumumab treatment. Data from *n* = 248 patients included in analysis. **B.** Average start date of daratumumab treatment in CoMMpass IA19 dataset = 46.8 +/-18.8 months (mean +/- S.D.). This timing of therapy appears consistent with divergence in overall survival curve (beginning at approximately month 45) between patients receiving daratumumab with tumors expressing top or bottom quintile of *CD38* transcript at initial diagnosis. Data from *n* = 80 patients included in analysis. *p*-value by log-rank test.

### Supplementary Methods

*Cell culture and cell line generation*

Development of RPMI-8226, AMO-1 and KMS12-PE multiple myeloma cell lines stably expressing the dCas9-KRAB machinery were described in our recent publication[4]. The FgH1tUTG plasmid (gift from Catherine Smith lab) containing the insert H1-Tet-sgRNA cassette for doxycycline inducible lentiviral guide RNA (Addgene 70183), was used to create the RPMI8226 dCas9-KRAB-FgH1tUTG/sgRNA cell line. MM.1S cells were originally obtained from ATCC; AMO-1 from DSMZ; KMS12-PE from JCRB. Cells were grown in RPMI-1640 complete growth media containing 10-20% fetal bovine serum (Gemini), 1% Penicillin/Streptomycin and 2 mM L-Glutamine. Cell lines were seeded at 0.2-0.4*$10^6$ cells/ml and passaged when they reached 1*$10^6$ cells/ml cell density. All cell lines were validated by STR profiling service provided by Genetica DNA Laboratories and tested for mycoplasma using Lonza MycoAlert kit.

*sgRNA libraries and CRISPRi screen*

The genome-wide CRISPRi v2.0 libraries subdivided into seven sublibraries containing five sgRNAs per gene were described previously[5]. For pooled screening, sgRNA libraries were packaged into lentivirus and harvested for transduction as described previously[5]. RPMI-8226 cells expressing the CRISPRi machinery were spin-infected with the virus at 700g for 2 hours at 32°C. Forty-eight hours later, the cells were analyzed for percentage of infection by using flow cytometry and were treated with 1 μg/mL of puromycin to obtain a pure population of single guide RNA (sgRNA)–expressing cells. On day 14 postinfection with the CRISPRi sublibraries, cells were stained for cell surface CD38 and flow-sorted to enrich for populations of cells expressing low or high cell surface levels of CD38. Briefly, for each sublibrary, cells were resuspended in fluorescence-activated cell sorter (FACS) buffer (phosphate-buffered saline containing 0.5% fetal bovine serum) at a concentration of 10 × $10^6$ cells/mL. The cells were blocked by using Human BD Fc Block (#564220; BD Biosciences), stained with FITC-CD38 antibody (BD Biosciences; clone HB7) and resuspended in FACS buffer for flow sorting. The top and bottom 25% of cells expressing CD38 as determined from FITC-CD38 histogram were flow-sorted by using FACSAria II (BD Biosciences). The various cell populations were then processed for next-generation sequencing as previously described[6] and sequenced on a HiSeq-4000 (Illumina). To identify significant hit genes, sequencing reads were analyzed by using the MAGeCK-iNC pipeline as previously described[7].

*Individual sgRNA knockdown*

Individual sgRNA sequences against specific targets were designed as described previously[7] and were cloned into lentiviral vector for expression from a U6 promoter[8]. Lentivirus production, transduction, and puromycin selection was performed for each individual guide RNA as was done for the library. Transduction was confirmed using flow cytometry. Knockdown of target gene was confirmed using flow cytometry or Western blotting.

*Doxycycline treatment*

For treatment of RPMI8226 dCas9-KRAB-FgH1tUTG/sgRNA cell line to induce expression of the sgRNA, doxycycline hyclate (Sigma-Aldrich D9891) was dissolved in sterile water at a stock concentration of 50 mg/ml as recommended per manufacturer, and added to RPMI-1640 media for a final concentration of 0.5 µg/ml, 1 µg/mL and 2 µg/mL for 72 hours.

*Flow cytometry*

Immunostaining of cells were performed as per the instructions from antibody vendor unless stated otherwise. Cells were pelleted and washed twice in FACS Buffer (PBS +2% FBS) before staining. Cells infected with CRISPRi libraries were resuspended at 5e6/ml and stained for 2 hours on ice. Cells for testing knockdowns or other test were resuspended at 1e6/ml and stained for 30 minutes on ice. Post-staining cells were washed twice with ice-cold FACS bufferSamples were immediately analyzed using either a BD Cytoflex or Sony SH800S or sorted on BD FACS Aria2 flow cytometer. Flow cytometry data was analyzed using FlowJo v10.8.1. Compensation when needed was performed with UltraComp eBeads - Compensation Beads (Thermo Fisher Scientific, 01-2222-42). Raw median fluorescence intensity (MFI) values of CD38 stained cells were normalized to isotype control samples and data plotted as fold change in MFI relative to untreated cells using Prism V7 (GraphPad Software) or RStudio v4.1.2 using tidyverse, readxl, reshape2 and ggpubr packages.

*Drug assays for surface CD38*

MM.1S and KMS11 cells were resuspended in fresh RPMI-1640 media and 2x10e5 cells were added to each well of a U-bottom 96-well plate. Cells were treated with increasing concentration of caffeic acid phenethyl ester (CAPE) (Millipore Sigma, C8221) or (*R*)-2-Hydroxyglutarate (Millipore Sigma, H8378) for 3 days after which CD38 expression was analyzed by flow cytometry (above).

*Western blotting*

Approximately $1x10^7$ cells were collected and washed with PBS three times, flash frozen, and stored at -80C prior to lysis and Western blot analysis. Cells pellets were lysed in 1X RIPA lysis buffer (Millipore 20-188) with HALT protease and phosphatase inhibitors (Thermo Fisher Scientific, 78442), incubated on ice for 15 minutes, sonicated for 15 seconds at 1 Hz cycles on ice, and cleared by centrifugation at 17000x*g* for 10 minutes at 4°C. Western blots were performed as previously described[9]. Primary antibodies for immunoblotting we used at manufacturer recommended dilutions: a-TLE3 (Santa Cruz, clone D-10), a-HEXIM1 (Santa Cruz, clone D-5Y5K, a-SPI1 (Cell Signaling Technology (CST), clone 2266), a-NFKB1 (CST, clone D4P4D), a-NFKB2 (CST, clone 18D10), a-MAPK (ERK1/2) (CST, clone 137F5), a-AKT (CST, clone 11E7), a-phospho-MAPK (ERK1/2)(Thr202/Tyr204)(CST, clone D13.14.4E), a-phospho-AKT (Ser473) (CST, clone D9E), a-XBP1 (Abcam 37152 ). Anti-b-actin (CST, clone 13E5) was used as loading controls for immunoblots. For XBP1 expression, the data was quantified in RStudio v4.1.2 using tidyverse, readxl, reshape2 and ggpubr packages.

*ADCC assays*

Assays were performed as in ref.[8]. Briefly, daratumumab was sourced from Janssen Pharmaceuticals. NK92-CD16 transgenic cells (a kind gift of Dr. Bruce Walchek, University of Minnesota) were then added an effector-to-target ratio of 20:1. After 20 hours, lysis was measured in a bioluminescence-based assay using CytoToxGlo (Promega). Lysis by ADCC was calculated using the following formula : % Lysis = (signal in presence of daratumumab – signal in presence of IgG1 control antibody) x100 / signal in presence of IgG1 control antibody.

*Murine studies*

Female NOD *scid* gamma (NSG; strain NOD.Cg-*Prkdc*$^{scid}$ *Il2rg*$^{tm1Wjl}$/SzJ) mice were obtained from Jackson Laboratories aged 6-8 weeks. RPMI-8226 CRISPRi cells were lentivirally transduced with a plasmid expressing luciferase and mCherry, as described previously[10]. 1e6 engineered RPMI cells (transduced with different guides) were I.V. injected into each NOD *scid* gamma mouse and were allowed to engraft for 14 days. Tumor burden was assessed by non-invasive bioluminescent imaging at the UCSF PTC on a Xenogen In Vivo Imaging System on days 14, 21, 28 and 35. 200ug of daratumumab was injected I.P. on Days 15, 17, 21, 24, and 28 post-implantation. No randomization was performed given different genetic alterations of tumor model in each arm. Sample size was not pre-determined based on power analysis and represented standard sample size for similar survival analysis in the field using this model. Investigators were not blinded to study conditions. All experiments were performed under approved protocols by the UCSF Institutional Animal Care & Use Committee (IACUC) and abided by all standard animal care and welfare guidelines.

*ChIP-seq and ATAC-seq analysis*

Raw data of ATAC-seq and H3K27Ac ChIP-seq data for Multiple Myeloma patients and normal Memory B cells, Plasmablasts, and Plasma cells were downloaded from European Nucleotide Archive database with the Data Accession Number: PRJEB25605 (ref.[1]). Paired-end reads were mapped to the human reference genome (hg19) using Bowtie2 for ChIP-seq and ATAC-seq reads. Only reads mapping uniquely to the genome with not more than 2 mismatches were retained for further analysis. Clonal reads (i.e., reads mapping at the same genomic position and on the same strand) were collapsed into a single read. Peaks from ChIPseq or ATAC-seq were called using the MACS2 (ref.[11] or Homer program[12]. We also checked the potential motif binding in the CD38 promoter regions. We predicted the motif binding in the identified ATAC peak regions from MM patients using PROMO tool[13], and version 8.3 of TRANSFAC.

*Patient gene expression microarray and RNA-seq data*

To identify transcription factors that may regulate CD38 expression, we calculated the gene expression correlation with CD38 expression of those identified transcription factors based on ATAC binding. Gene expression RNAseq or microarray data from three large cohorts of patients with MM were analyzed, including Multiple Myeloma Research Foundation CoMMpass dataset (*n* = 664, IA13, https://research.themmrf.org/), the MAQC-II Project: Multiple myeloma (MM) data set (*n* = 554, GSE24080 (ref.[14]), and the MM clinical trials of bortezomib (*n* = 264, GSE9782 (ref.[15]). Expression level of a gene in a sample was determined by the average of expression values from multiple probesets on the array representing this gene or form the FPKM values of RNA-seq. The correlation p-values of expression value of two genes were determined by two-sided Spearman and Pearson correlation. All downstream microarray analysis was performed using R version 2.14.0.

*Machine learning analysis for transcriptional determinants of CD38 expression*

Patient transcriptomic data was acquired from the Multiple Myeloma Research Foundation CoMMpass dataset (IA13; https://research.themmrf.org/). A list of 100 transcription factors was generated for this analysis by combining those from ATAC-seq motif analysis of MM patient tumors (above) with analysis of ENCODE data suggesting binding of different transcription factors to the *CD38* locus in non-plasma cells. CD138+ enriched RNA-Seq data from 779 newly-diagnosed patients was incorporated into the model. All counts (in Transcripts per Million (TPM)) were log-transformed prior to model building and analysis.

We developed an XGBoost model (Extreme Gradient Boosting) using the xgboost package. The code for our analysis is available at our repository here (https://github.com/vsarin92/CD38_Predictor) and the environment yaml file can be used to recreate the specific anaconda environment (with all the relevant packages) to build the models and run the analysis themselves.

Initially all transcription factors determined to be co-expressed were utilized together to build the model. We used randomized search with cross validation to find the best parameters for the XGBoost models to predict *CD38* expression. Specifically we conducted a search over colsample_bytree (subsample ratio of features when constructing each tree), colsample_bylevel (subsample ratio of features for each level), colsample_bynode (subsample ratio of features for each split), gamma (minimum loss reduction to partition a leaf node), learning_rate (how big of a step to take after each iteration), max_depth (maximum tree depth), n_estimators (number of trees), and subsample (subsample ratio of training data). Random combinations of these were used to build a model with 10-fold cross validation, for 50000 iterations, and the hyperparameters that gave the highest cross validation score was what was utilized in further model building.

We conducted SHAP analysis on each model built to help describe what affect features have on *CD38* model predictions using the shap package. Essentially for each patient, a feature is selected to be replaced by a random value from the distribution for that feature. The model's prediction for this simulated patient is compared with the average predictions of all other patients and the difference is attributed to the randomly selected feature. This is repeated for each patient and each feature. Note that SHAP analysis was conducted only with test data (20% of CoMMpass dataset).

Model performance was validated by calculating coefficient of determination ($R^2$) and adjusted $R^2$ values (which accounts for the number of observations and the number of features used). Furthermore, the model underwent 5-fold cross validation with the mean absolute error, $R^2$, and adjusted $R^2$ being reported for each fold to validate that performance was not dependent on our original choice of training and test data.

*Drug treatments for mass spectrometry samples*

Multiple myeloma cell lines at $5e10^5$ cells/mL were treated with 10 nM ATRA (Sigma) and 10 nM panobinostat (Selleck chemicals) for three days continuously. Multiple myeloma cells were treated with addition of freshly prepared azacytidine (Sigma) in fresh media every day of the treatment. Cells were treated continuously for three days (3d) and allowed to recover for four days in fresh media as described previously[8]. Post drug-treatment, live cells were enriched using Histopaque density-gradient centrifugation before cell surface protein labeling.

*Cell surface protein labeling*

Cell surface proteins were labeled with biotin using the N-linked glycosylation-site biotin labeling method[16]. Briefly, $3e10^7$ live cells were washed twice and resuspended in 1 mL of ice-cold PBS, and treated with 1.6mM sodium metaperiodate (VWR, 13798-22) at 4°C for 20 minutes to oxidize the vicinal diols of sugar residues linked to surface proteins. The cells were then washed twice in PBS in order to remove excess sodium metaperiodate. Cells were resuspended in 1 mL of ice-cold PBS and treated with 1 mM biocytin hydrazide (Biotium, 90060) and 10 mM aniline (Sigma-Aldrich, 242284) at 4°C for 90 minutes with gentle mixing in order to biotinylate free aldehydes exposed on the sugar residues. After labeling, cells were washed three times with ice-cold PBS to removed excess biotin, frozen in liquid nitrogen, and stored at -80°C until further processing for mass spectrometry. All experiments were performed in biological triplicate with replicates harvested from consecutive passages.

*Cell lysis, cell surface protein enrichment, and peptide digestion*
Frozen cell pellets were thawed on ice in 1ml of RIPA buffer (Millipore, 20-188) with the addition of 1X 22 HALT protease inhibitors (Pierce, 78442). After incubation on ice for 10 minutes, cells were disrupted by sonication and the lysates were clarified by centrifugation at 17,000 RCF at 4°C for 10 minutes. Clarified lysate was mixed with 500 uL of neutravidin agarose resin (Thermo, 29200) and incubated at 4°C for 2 hours with end-over-end mixing. Neutravidin beads with captured biotinylated surface proteins were washed extensively by gravity flow to remove unbound proteins using 50 mL of 1X RIPA + 1mM EDTA, followed by 50 mL of PBS + 1M NaCl, and finally 50 mL of 50 mM ABC + 2M Urea buffer. Washed beads were resuspended in digestion buffer (50 mM Tris pH 8.5, 10 mM TCEP, 20 mM 2-Iodoacetamide, 1.6M Urea) with 10ug of added Trypsin protease (Pierce, 90057) to perform simultaneous disulfide reduction, alkylation and on-bead peptide digestion at room temperature overnight (16-20 hours). After digestion, the pH was dropped to ~2 with neat trifluoroacetic acid (TFA, Sigma, T6508-10AMP) and the peptide mixture was desalted using a SOLA-HRP column (Thermo, 60109-001) on a vacuum manifold. Desalted peptides were eluted with 50% acetonitrile (ACN, Sigma, 34998-4L) and 50% water with 0.1% TFA and dried down completely in a speedvac. Dried peptides were resuspended in LC/MS grade water (Fisher, W64) with 2% 10 ACN and 0.1% formic acid (FA, Honeywell, 94318-250ML-F). Peptide concentration was measured using 280 nm absorbance on a Nanodrop (Thermo), and the peptide concentration was adjusted to 0.2ug/ul for mass spec runs.

*Surface proteomics LC-MS and Data Analysis*
For each replicate, 1 ug of peptide was injected onto a Dionex Ultimate 3000 NanoRSLC instrument with a 15-cm Acclaim PEPMAP C18 (Thermo, 164534) reverse phase column. The samples were separated on a 3.5-hour non-linear gradient using a mixture of Buffer A (0.1% FA) and B (80% ACN/0.1% FA), from 2.4% ACN to 32% ACN. Eluted peptides were analyzed with a Thermo Q-Exactive Plus mass spectrometer. The MS survey scan was performed over a mass range of 350-1500 m/z with a resolution of 70,000, with a maximum injection time (IT) of 100 ms. We performed a data-dependent MS2 acquisition at a resolution of 17,500, AGC of 5e4, and IT of 150 ms. The 15 most intense precursor ions were fragmented in the HCD at a normalized collision energy of 27. Dynamic exclusion was set to 20 seconds to avoid over-sampling of highly abundant species. The raw spectral data files have been deposited at the ProteomeXchange PRIDE repository (accession number PXD027594).

Raw spectral data was analyzed using MaxQuant v1.5.1.2[17] to identify and quantify peptide abundance and searched against the human Swiss-Prot annotated human proteome from Uniprot (downloaded 5/13/18 with 20,303 entries). The "match-between-runs" option was selected to increase peptide identifications while the "fast LFQ" option was selected to calculate label-free quantification values (LFQ) of identified proteins. All other settings were left to the default MaxQuant values. The MaxQuant output data was analyzed using Perseus[18] and the R program (version 3.4.0) in R-Studio. Proteins annotated as "reverse", "only identified by site", and "potential contaminant" were filtered out as well as proteins that were quantified in less than 2 out of 3 biological replicates in at least one experimental group. Proteins were further filtered to include only membrane-proteins or membrane-associated proteins using a manually curated list of surfaceome proteins[19]. Missing values were imputed based on the normal distribution of the dataset as implemented by Perseus. Volcano plots were generated using output from a two-sample *t*-test comparing the $\log_2$ transformed LFQ protein abundance values from different cell lines with a false discovery rate (FDR) set to 0.01.

*RNA-seq*
RNA-sequencing and data analysis RNA extraction, library preparation, and sequencing were performed at BGI (Shenzhen, China). Samples were sequenced using the BGISEQ-500 platform, with a minimum of 21.4 million reads per sample. For genome mapping, clean reads were mapped to reference genome using HISAT2 with an average of 62% unique read mapping across samples. For gene expression analysis, clean reads were mapped to reference transcripts using Bowtie2 and expression levels calculated using RSEM. Transcripts per million (TPM) were used for data analysis performed in *R*. Raw sequencing data has been deposited at the Gene Expression Omnibus (GEO) (accession number GSE181277).

*Phosphoproteomics sample preparation*
RPMI-8266 cells were serum-starved in RPMI-1640 media containing 1% FBS for 22-24 hours at 1e6/mL. Control IgG or Daratumumab was added for a final concentration of 20 mg/ml and allowed to bind for 20 minutes. Immediately after, the media (containing cells) was transferred to pre-chilled tubes, pre-chilled low-serum media was added to the plates and adherent cells were scraped while on ice. Cells were pelleted from collected media, washed twice with ice-cold PBS, flash-frozen, and stored in -80°C before processing. Triplicates of each condition were obtained. The same conditions were used for confirmatory Western blotting at the noted time points (see antibody details above).

We lysed the samples in 6M guanidine hydrochloride (GdnHCl), 0.1M Tris pH 8.5, 5 mM TCEP, 10 mM 2-chloroacetamide and sonicated the lysate at 1 Hz pulses for 45 seconds on ice. Protein concentrations were quantified using the 660nm Protein Assay (Pierce 22660). For each sample, 1 mg of protein was transferred into a clean tube and diluted to 1M GdnHCl by 0.1M Tris pH8.5. Next, we added 20 ug of MS-grade trypsin (Thermo, PI90057) and incubated the mixture for 20 hours at 37°C.

We halted the trypsin digestion by acidification with trifluoroacetic acid (TFA) to 1% (vol/vol). Any precipitate was removed by centrifugation at 17,200*g* for 5 minutes. The samples were then desalted on a SOLA C18 cartridge (Thermo, 03150391) assisted by vacuum. We washed twice with 0.1% TFA followed by another 2% acetonitrile (ACN)/0.1% formic acid (FA) wash. We eluted the peptides in 80% ACN/0.1% TFA.

Next, we enriched for phospho-peptides using a $Fe^{3+}$-immobilized metal affinity column (IMAC). We prepared the column by stripping the nickel off Ni-charged agarose beads (VWR, 220006-720) using four washes of 100mM EDTA and re-charging them with $Fe^{3+}$ from 150mM $FeCl_3$. We then transferred the $Fe^{3+}$-beads to a MicroSpin C18 column (Nest Group, SEM SS18V.25). After IMAC preparation, we transferred our tryptic peptides to the column and performed three washes, under vacuum, with 80% ACN/0.1% TFA to remove non-phosphorylated peptides and two washes with 0.5% FA to equilibrate the C18 resin. To mobilize the phospho-peptides onto C18, we washed twice with 500mM potassium phosphate at pH 7. We then performed two final washes with 0.5% FA to eliminate any residual salt. The phospho-peptides were eluted in 50% ACN/0.1% FA, vacuum dried, and stored at -80°C for further analysis.

*Phosphoproteomic liquid chromatography-tandem mass spectrometry (LC-MS/MS) analysis*
Enriched phospho-peptides were re-suspended in 2% ACN/0.1% FA. A total of 1 ug of peptides from each sample were injected into a Dionex Ultimate 3000 NanoRSLC instrument with a 15-cm Acclaim PEPMAP C18 (Thermo, 164534) reverse phase column. The samples were separated on a 3.5-hour non-linear gradient using a mixture of Buffer A (0.1% FA) and B (80% ACN/0.1% FA). The initial flow rate was 0.5 uL/min at 3% B for 15 minutes followed by a drop in flow rate to 0.2 uL/min and a non-linear increase (curve 7) to 40% B for the next 195 minutes. The flow rate was then increased to 0.5 uL/min while Buffer B was linearly ramped up to 99% for the next six minutes. Finally, we maintained the peak flow rate and Buffer B concentration for another seven minutes before dropping the concentration back to 3%.

Eluted peptides were analyzed with a Thermo Q-Exactive Plus mass spectrometer. The MS survey scan was performed over a mass range of 350-1500 m/z with a resolution of 70,000. The automatic gain control (AGC) was set to 3e6, and the maximum injection time (MIT) was 100 ms. We performed a data-dependent MS2 acquisition at a resolution of 17,500, AGC of 5e4, and MIT of 150 ms. The 15 most intense precursor ions were fragmented in the HCD at a normalized collision energy of 27. Dynamic exclusion was set to 20 seconds. The raw spectral data files are available at the ProteomXchange PRIDE repository (Accession number PXD027594).

*Phosphoproteomics data processing*
We analyzed the raw spectral data using MaxQuant (version 1.5.1.2) to identify and quantify phospho-sites on serine, threonine, and tyrosine. We relied on default settings except for the inclusion of “Phospho (STY)” as a variable modification. We searched against the Swiss-Prot-annotated human proteome from UniProt (downloaded August 2016 with 20,163 entries).
We processed the “Phospho (STY) Sites” output file containing the phospho-sites data for downstream analyses in R (version 3.6.3). First, we filtered out proteins annotated as “Reverse” or “Potential contaminant” and retained sites with “localization probability” and “delta score” greater than 0.75 and 8, respectively. To remove poorly quantified phospho-sites, we required each site to be quantified in at least two replicates in one condition. This reduced the number of identifications from 6,317 to 5,430. We then $\log_2$-transformed the intensity data and centered the median intensity of each sample at 0 by subtracting the sample median from each value. Missing values were imputed using a hybrid imputation approach, “MLE” setting for values missing at random and “MinProb” for those missing not at random (*imputeLCMD* package (https://cran.r-project.org/web/packages/imputeLCMD/imputeLCMD.pdf)). Next, we calculated differential phosphorylation using Welch’s unpaired two-sample *t*-test with Benjamini-Hochberg correction for determining false discovery rate. Additionally, PTM Signature Enrichment Analysis (https://github.com/broadinstitute/ssGSEA2.0) was performed to extract differentially regulated signaling pathways in the daratumumab-treated condition compared to the IgG-treated control.

*Kinase-Substrate Enrichment Analysis*

We implemented the Kinase-Substrate Enrichment Analysis (KSEA) on our phospho-proteomic data in *R*. First, we acquired kinase-substrate associations from a manually curated database called PhosphoSitePlus (downloaded 2/5/2018 with 10,188 human entries) and from a recent publication on the large-scale identification of *in vitro* kinase substrates[20]. We combined the kinase-substrate associations from PhosphoSitePlus and from the publication to obtain 190,336 interactions spanning 27,652 phospho-sites and 430 kinases. Using these annotations, we then inferred kinase activities by averaging the intensities of phospho-sites associated with a given kinase. The *p*-value of the kinase activity score was determined using Welch's one-sample *t*-test on the log2FCs of kinase substrates followed by Benjamini-Hochberg correction for multiple hypothesis testing.

*Data availability*

RNA-seq: Raw sequencing data has been deposited at the Gene Expression Omnibus (GEO) (accession number GSE181277).
Proteomics: The raw spectral data files are available at the ProteomeXchange PRIDE repository (Accession number PXD027594).

*Statistical analysis:*

All data is presented as mean +/- S.D. or S.E.M. as described in the associated figure legend. No samples were specifically excluded from analyses. Sample sizes were not pre-determined to give a specific effect size. Investigators were not blinded to study conditions. Statistical significance in proteomics and transcriptomics analysis was determined by Welch's *t*-test; two-sample *t*-test with null hypothesis that the different in $\log_2$-transformation of proteomic LFQ's, transcriptome FPKM's for RNA-seq, or transcriptome microarray intensity is equal 1 to 0. An unadjusted $p < 0.05$ is considered statistically significant. Number of biological replicates is specified in associated figure legend. For *in vitro* assays number of biological and technical replicates are specified within the associated figure legend. *p*-values were obtained for *in vitro* assays by two-sided *t*-test or one-way ANOVA as specified in the associated figure legend. Variance was estimated to be similar between compared groups and appropriate statistical tests used. Correlation plots display Pearson correlation *R* and associated *p*-value by two-sided *t*-test. Throughout the manuscript for p-values **** <0.001, ***<0.005, **<0.01, *<0.05.

**Supplementary Table Legends (Tables attached as Excel files)**

**Supplementary Table 1. Results of CRISPRi screen.** Data display average fold-change in CD38 integrated across 5 sgRNA's/gene ("epsilon" value) as determined by MAGeCK pipeline. Positive epsilon indicates gene knockdown led to increased surface CD38. "NTC" indicates non-targeted control sgRNA.

**Supplementary Table 2. ATAC-seq analysis.** Transcription factors identified by ATAC-seq binding peaks and PROMO motif prediction (Columns I-R), and their expression correlation with CD38 expression in three MM clinical trials (Columns A-G and T-AP). Columns A-G are the summary of Pearson correlation and Spearman correlation from the three clinical trials. Columns I-R are the transcription factors and their presence (1) or absence (0) in each of the eight peaks in the CD38 promoter region identified by ATAC-seq binding peaks and PROMO motif prediction tool. Columns T-AP are the detailed correlation coefficient and p values from Pearson or Spearman correlation test for each of the three clinical trials.

**Supplementary Table 3. CD38 "antigen escape profiling".** *Sheet 1.* MaxQuant + Perseus outputs of cell surface proteomic analysis aggregated across RPMI-8226, AMO-1, and KMS12-PE cells after CD38 knockdown by CRISPRi, compared to non-targeting sgRNA (scri). *Sheet 2.* Integrated summary outputs of cell surface proteomic analysis fold-change with RNA-seq after CD38 knockdown across all cell lines.

**Supplementary Table 4. Proteomic impact of drug treatment.** *Sheets 1-3.* MaxQuant + Perseus outputs from cell surface proteomics after RPMI-8226 treatment by Aza, ATRA, or Pano. *Sheets 4-5.* Integrated final outputs for fold-change with RNA-seq for Aza and ATRA.

**Supplementary Table 5. Phosphoproteomics after daratumumab.** *Sheet 1*. MaxQuant + Perseus outputs from phosphoproteomics after daratumumab treatment of RPMI-8226. *Sheet 2*. Results of KSEA analysis from analyzed phosphopeptides.
